# Long-term ecological and evolutionary dynamics in the gut microbiomes of carbapenemase-producing *Enterobacteriaceae* colonized subjects

**DOI:** 10.1101/2022.05.11.491472

**Authors:** Jonathan T. L. Kang, Jonathan Teo, Denis Bertrand, Amanda Ng, Aarthi Ravikrishnan, Melvin Yong, Ng Oon Tek, Kalisvar Marimuthu, Swaine Chen, Chng Kern Rei, Gan Yunn-Hwen, Niranjan Nagarajan

## Abstract

Long-term colonization of the gut microbiome by carbapenemase-producing *Enterobacteriaceae* (CPE) is a growing area of public health concern as it can lead to community transmission and rapid increase in cases of life-threatening CPE infections. Leveraging the observation that many subjects are decolonized without interventions within a year, we used longitudinal shotgun metagenomics (up to 12 timepoints) for detailed characterization of ecological and evolutionary dynamics in the gut microbiome of a cohort of CPE-colonized subjects and family members (n=46; 361 samples). Subjects who underwent decolonization exhibited a distinct ecological shift marked by recovery of microbial diversity, key commensals and anti-inflammatory pathways. In addition, colonization was marked by elevated but unstable *Enterobacteriaceae* abundances, which exhibited distinct strain-level dynamics for different species (*Escherichia coli* and *Klebsiella pneumoniae*). Finally, comparative analysis with whole genome sequencing data from CPE isolates (n=159) helped identify sub-strain variation in key functional genes and the presence of highly similar *E. coli* and *K. pneumoniae* strains with variable resistance profiles and plasmid sharing. These results provide an enhanced view into how colonization by multi-drug resistant bacteria associates with altered gut ecology and can enable transfer of resistance genes, even in the absence of overt infection and antibiotic usage.

## Introduction

The global dissemination of antibiotic resistance genes among pathogenic bacteria is a major public health problem that, if left unaddressed, would lead to reduced efficacy of current treatment options, elevated treatment costs, and increased mortality^1^. A particular area of concern is the spread of carbapenemase-producing *Enterobacteriaceae* (CPE)^2,3^, with their ability to degrade carbapenems often acquired in gram-negative bacteria from plasmids with carbapenemase genes^4,5^, thus rapidly endangering the utility of these antibiotics of last resort^6,7^. In addition to causing life-threatening infections, asymptomatic colonization of CPE in the human gut is increasingly common^8,9^, creating a reservoir for transmission of antibiotic resistance^10^. While prior CPE studies have focused on epidemiology^3,11^ and molecular aspects^7,12,13^, the natural history of gut colonization including ecological and evolutionary changes linked to antibiotic resistance transmission or CPE decolonization remain unexplored.

In recent years, studies into host-microbiome-pathogen interactions have provided important insights into pathogenesis^14^, immune response^15^ and treatment avenues^16^ for various viral and microbial pathogens. These studies typically leverage metagenomic approaches to track microbial community composition over time and understand ecological responses to overt infection^16,17^. As microbial populations often have rapid turnover, whole-genome sequencing of pathogenic isolates has been used to study intra-host evolution during chronic infections, identifying key enzymes for host adaptation and colonization^18,19^. Alternatively, deep shotgun metagenomic sequencing can simultaneously reveal nucleotide level variation for many bacterial species of interest^20,21^, shedding light on strain-level dynamics in the community. This approach has been used to study stable microbiomes in healthy individuals as well as dynamic changes during fecal microbiota transplantation^22^. Asymptomatic gut colonization of CPE strains presents a unique opportunity to study an intermediate phenomenon i.e. strain competition with commensals, and associated ecological and evolutionary adaptations, in the absence of an overt infection or disease.

Here we conducted longitudinal gut microbial analysis for a cohort of index subjects (n=29, CPE colonized at recruitment) and their family members (n=17, not CPE colonized) with up to 12 time points over the duration of a year, to obtain multiscale^23^ (microbiome composition, strains and gene-level) characterization of ecological and evolutionary changes during CPE colonization. Based on deep shotgun metagenomic sequencing of stool DNA, we observed distinct ecological shifts marked by recovery of diversity and key commensals in association with CPE decolonization. CPE colonization was marked by elevated but unstable *Enterobacteriaceae* abundances, which exhibited specific dynamics at the strain-level for different species (*Escherichia coli* and *Klebsiella pneumoniae*). Comparative analysis with whole genome sequencing data from CPE isolates (n=159) helped identify the presence of highly similar *E. coli* and *K. pneumoniae* strains with variable resistance profiles and plasmid sharing. These results provide an enhanced view into how colonization by multi-drug resistant bacteria associates with altered gut ecology and can enable transfer of resistance genes, even in the absence of overt infection and antibiotic usage.

## Results

### CPE colonization is associated with ecological shifts that are resolved during recovery

Leveraging the observation that CPE carriage in hospital patients can be resolved within 3 months, with 98.5% probability within a year for our cohort^24^ (though other cohorts have reported longer durations^25,26^), we tracked gut microbiome composition in this cohort of individuals for a year to understand ecological changes associated with decolonization (up to 12 timepoints, 361 samples in total; **Table 1, Supplementary File 1**). Specifically, stool samples were obtained from hospital patients who screened positive for CPE carriage (n=29, index subjects), as well as their non-CPE-colonized family members (n=17, serving as home environment-matched controls) and characterized via deep shotgun metagenomic sequencing (>50 million Illumina 2×100bp reads, on average; **Methods**). Principal coordinates analysis with average-linkage clustering based on taxonomic profiling of the data showed that there are multiple distinct community configurations (I, II, III, IV), where CPE positive samples (based on stool culture and qPCR^24^) were less commonly seen in configurations I and II, and more commonly seen in configurations III and IV (**Figure 1a, Supplementary Figure 1, Supplementary File 2**). This statistically significant shift of CPE positive samples along PCoA1 (Wilcoxon rank-sum p-value<1.4×10^−8^, **Supplementary Figure 2a**) is defined by a gradient of relative abundances that are most strongly correlated for the genera *Escherichia* (negative i.e. more abundant in configuration IV samples) and *Bacteroides* (positive; **Supplementary Figure 2b**). A similar shift was observed when comparing taxonomic profiles for configuration IV versus configuration I microbiomes (**Supplementary File 3**). Interestingly, while configuration IV has no microbiomes from family members, a few CPE negative samples from index subjects also cluster here.

**Table 1.** Cohort and sample statistics. Stool samples were collected at recruitment (V00), weekly for 4 weeks (V01-V04), monthly for 5 months (V05-V09) and bimonthly for 6 months (V10-V12) when provided by study participants.

|  | Number of<br>individuals | Total number of<br>samples | Mean number of samples<br>per individual |
| --- | --- | --- | --- |
| **Index subjects** | 29 | 216 | 7.5 |
| **Family members** | 17 | 145 | 8.5 |
| **Total** | 46 | 361 | 7.9 |

**Figure 1.**
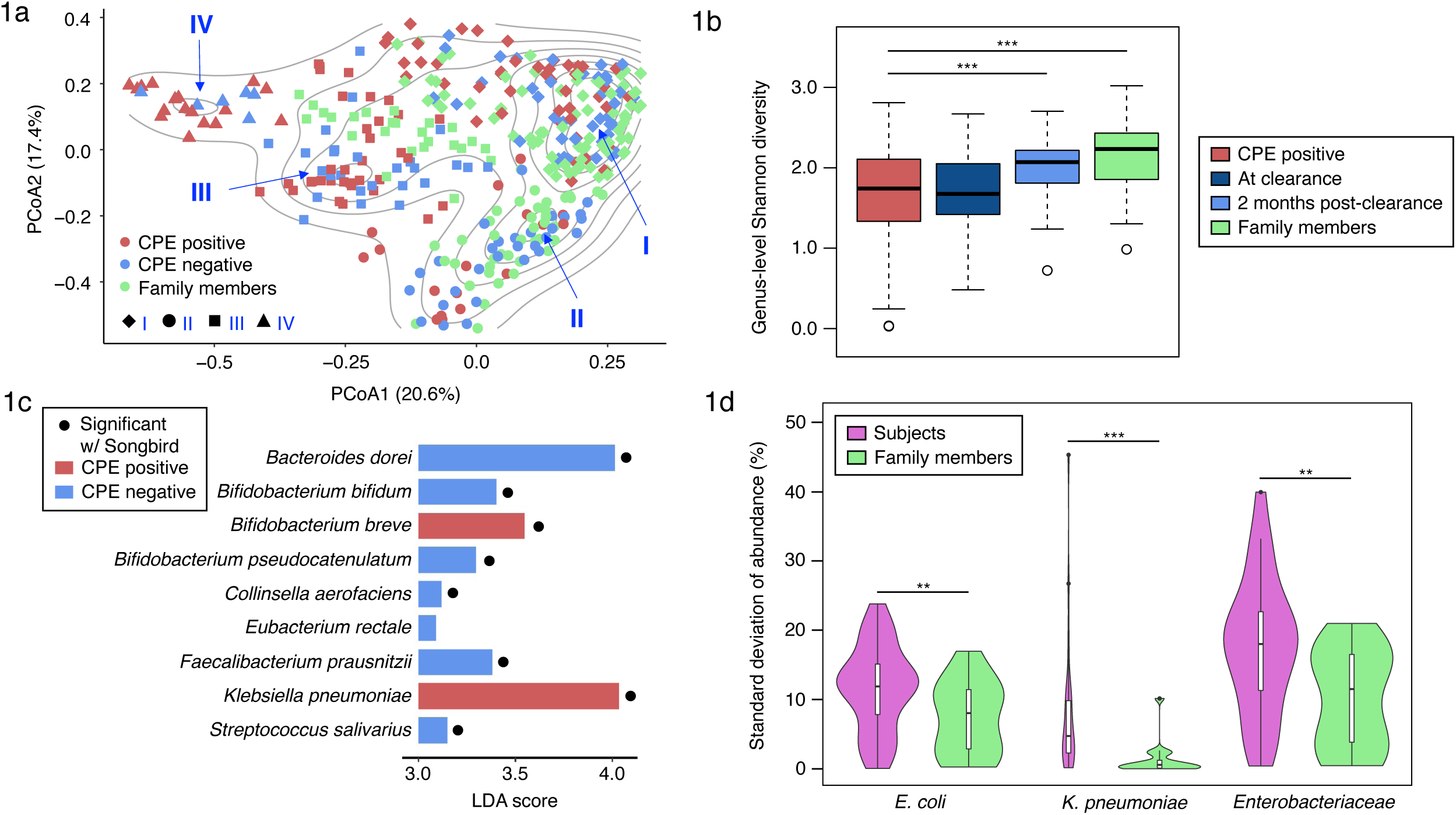
Shifts in gut microbial ecology associated with CPE colonization. (a) Principle coordinates analysis plot showing how gut microbial community composition varies in relation to CPE colonization status (genus-level Bray-Curtis dissimilarity; Unifrac plot in **Supplementary Figure 1**). Contour lines indicate similar density with regions of locally higher density associated with distinct community configurations (defined by average linkage clustering in **Supplementary Figure 1**; marked with labels I, II, III and IV). (b) Boxplots showing genus-level Shannon diversity distributions for different timepoints for index patients (“CPE positive” = during colonization, “At clearance” = within 1 month of decolonization, and “2 months post-clearance” = time points that were >2 months after decolonization), and all timepoints for family members. (c) Species that were found to be enriched in CPE positive (in red) and CPE negative (in blue) samples along with their LDA scores based on LEfSe analysis. Results that were significant based on Songbird analysis as well are indicated with a solid circle. (d) Violin plots showing the standard deviation of relative abundances (ignoring relative abundances <0.1% to avoid the effect of detection limit for metagenomics) over time of various taxa in different individuals (subjects and family members). For all subfigures * = p<0.1, ** = p<0.05, *** = p<0.01 based on the Wilcoxon rank-sum test and all other comparisons were not statistically significant.

Grouping timepoints based on their proximity in time to CPE clearance, highlighted that while CPE positive samples have the lowest average diversity^27^, there is a gradual increase in diversity around the time of decolonization and post decolonization, with diversity reaching the higher levels seen in family members after 2 months (**Figure 1b**). This pattern was seen even after accounting for potential confounding factors including antibiotic usage, hospitalization status, multiple timepoints for an individual, gender and ethnicity in a linear mixed-effects model (**Supplementary Figure 3**; **Methods**). We investigated if colonization of *Enterobacteriaceae* species alone could explain these changes by computationally subtracting all of them from taxonomic profiles and recomputing diversity metrics. We noted that both genus-level richness and Shannon diversity consistently preserved the trend of increasing during and after decolonization (**Supplementary Figure 4**), suggesting that these observations do not have a simplistic explanation due to CPE colonization, and point to a more pronounced shift in the microbiome.

The temporal shifts in diversity during CPE colonization were also reflected in terms of overall similarity among microbiomes, with Bray-Curtis distances (genus-level) to family members being highest in CPE positive samples, gradually reducing during and post de-colonization towards baseline values seen among family members (**Supplementary Figure 5**). These results highlight the ecological shift associated with CPE colonization that largely resolves post decolonization, but might have residual effects in some individuals.

**Table 2.** Top genes with sub-strain variation. Top 10 genes with most function-altering SNVs (≥10) with low frequencies (AF<0.5) for *E. coli* and *K. pneumoniae*. SNVs in distinct individuals were counted separately, but multiple timepoints were counted as one. P-values were computed by performing a binomial test on the observed number of SNVs given the probability of a gene acquiring SNVs after accounting for 1) the probabilities of acquiring mutations across all genes and 2) the codon composition of the genes where the respective SNVs are found. Bonferroni correction (α=0.05) was applied.

| Gene | Protein | SNV count | P-value |
| --- | --- | --- | --- |
| *E. coli* |  |  |  |
| ECIAI39_4258 | Putative invasin/intimin protein | 48 | $<10^{-10}$ |
| ECIAI39_0530 | Host specificity protein J of prophage | 34 | $<10^{-13}$ |
| ydcM | Putative transposase | 24 | $<10^{-15}$ |
| ECIAI39_1027 | Putative GTP-binding domain | 19 | $<10^{-12}$ |
| ECIAI39_4240 | Putative antirestriction protein | 18 | $<10^{-14}$ |
| yeeS | Putative DNA repair protein; CP4-44 prophage | 18 | $<10^{-16}$ |
| lacZ | Beta-D-galactosidase | 17 | $<0.05$ |
| ECIAI39_4893 | Putative tail fiber component K of prophage | 14 | $<10^{-7}$ |
| lacY | Lactose/galactose transporter | 14 | $<10^{-4}$ |
| Rz | Endopeptidase from phage origin (Lysis protein) | 13 | $<10^{-9}$ |
| ECIAI39_1018 | Hypothetical protein | 11 | $<10^{-4}$ |
| ECIAI39_4867 | Hypothetical protein | 10 | $<10^{-2}$ |
| yncK | Transposase ORF A | 10 | $<10^{-8}$ |
| *K. pneumoniae* |  |  |  |
| KPHS_51490 | Putative transposase | 30 | $<10^{-39}$ |
| KPHS_35720 | Putative transposase | 18 | $<10^{-13}$ |
| KPHS_22580 | Hypothetical protein | 12 | $<10^{-4}$ |

**Table 3.** Genes containing non-synonymous SNVs that exhibit a large (>0.3) change in allele frequency across pairs of “one strain” time points in an individual, observed across at least two individuals.

| Gene | Protein | SNV position<br>(no. of indiv.) | Ref |
| --- | --- | --- | --- |
| *E. coli* |  |  |  |
| **lacZ** | Beta-D-galactosidase | Cys77Arg (2),<br>Val86Ile (2),<br>Thr109Ala (2),<br>Phe114Tyr (2),<br>Ala117Thr (3),<br>Ser133Cys (2),<br>Leu350Val (2),<br>Ser656Thr (2),<br>Ser656Leu (2),<br>Thr746Met (2),<br>Asp753Gly (3),<br>Ala821Val (2) | Ref<sup>83</sup> |
| **pflB** | Pyruvate formate lyase I | Thr16Ala (3),<br>Asn383Ser (3) |  |
| **lpp** | Murein lipoprotein | Ala3Pro (2),<br>Lys2Asn (3) | Ref<sup>84</sup> |
| **pnp** | Polynucleotide<br>phosphorylase/polyadenylase | Ile122Val (3) | Ref<sup>43</sup> |
| **rnpA** | Protein C5 component of RNase P | Lys91Arg (3) |  |
| **rpsE** | 30S ribosomal subunit protein S5 | His121Arg (3) |  |
| **pheT** | Phenylalanine tRNA synthetase,<br>beta subunit | Glu699Asp (3) |  |
| **rsxC** | Putative 4Fe-4S ferredoxin-type<br>protein fused with unknown protein | Val712Ile (3) |  |
| **srmB** | ATP-dependent RNA helicase | Ser14Asn (3) | Ref<sup>85</sup> |
| **thrS** | Threonyl-tRNA synthetase | Lys638Gln (3) | Ref<sup>86</sup> |
| **yfgA** | Conserved hypothetical protein;<br>putative HTH-type transcriptional<br>regulator | Gln332Pro (3) |  |
| **nuoE** | NADH:ubiquinone oxidoreductase, chain E | Met26Lys (3) | Ref<sup>45</sup> |
| **nlpl** | Lipoprotein precursor | Thr57Ser (3) |  |
| **atpB** | F0 sector of membrane-bound ATP synthase, subunit a | Ile236Val (3) |  |
| **dnaK** | Chaperone Hsp70, co-chaperone with DnaJ | Ala449Ser (3) | Ref<sup>87</sup> |
| **eno** | Enolase | Glu132Ala (3) | Ref<sup>88</sup> |
| **gapA** | Glyceraldehyde-3-phosphate dehydrogenase A | Ile66Val (3) |  |
| **holC** | DNA polymerase III, chi subunit | Val136Met (3) |  |
| ***K. pneumoniae*** |  |  |  |
| **fadB** | 3-hydroxyacyl-CoA dehydrogenase/3-hydroxybutyryl-CoA epimerase/delta(3)-cis-delta(2)-trans-enoyl-CoA isomerase/enoyl-CoA hydratase | Ala725Asp (2) |  |
| **srmB** | ATP-dependent RNA helicase | Asn14Ser (2) | Ref<sup>85</sup> |
| **KPHS_52800** | Ribonuclease P protein component | Arg9Lys (2) |  |
| **KPHS_48690** | Hypothetical protein | Leu28Met (2) |  |
| **rpoA** | DNA-directed RNA polymerase subunit alpha | Ala157Thr (2) | Ref<sup>89</sup> |
| **rpIM** | 50S ribosomal protein L13 | Glu86Gln (2) |  |
| **KPHS_47160** | Hypothetical protein | Val10Ile (2) |  |
| **pnp** | Polynucleotide phosphorylase/polyadenylase | Val99Ile (2) | Ref<sup>43</sup> |
| **nlpl** | Lipoprotein | Ser57Thr (2) |  |
| **deaD** | ATP-dependent RNA helicase | Thr22Asn (2) |  |
| **rodZ** | Cytoskeleton Protein | Ser327Pro (2) |  |
| **lpxD** | UDP-3-O-[3-hydroxymyristoyl] glucosamine N-acyltransferase | Leu9Phe (2) | Ref<sup>90</sup> |
| **KPHS_36220** | Hypothetical protein | Val20Met (2) |  |
| **pheT** | Phenylalanyl-tRNA synthetase subunit beta | Asp699Glu (2) |  |
| **KPHS_27510** | Hypothetical protein | Ser29Tyr (2) |  |
| **gapDH** | Glyceraldehyde-3-phosphate dehydrogenase | Val66Ile (2) |  |
| **loID** | Lipoprotein-releasing system ATP-binding protein | Asn19Ser (2) |  |
| **mukE** | Condesin subunit E | Leu3Ser (2) |  |
| **KPHS_15690** | Alpha-ketoglutarate decarboxylase | Asp935Glu (2) |  |
| **KPHS_15680** | succinate dehydrogenase iron-sulfur subunit | Lys2Arg (2) |  |

To further probe into key bacterial species associated with CPE colonization we conducted differential abundance analysis based on CPE status (**Methods, Supplementary File 2**). While most *Enterobacteriaceae* species were not differentially abundant, *Klebsiella pneumoniae*^3,7^ had one of the strongest associations with CPE positive status (**Figure 1c**). In addition, only one other species (*Bifidobacterium breve*) was significantly enriched in CPE positive samples, while 7 other species were significantly depleted relative to CPE negative samples. These included several important commensal species that are known to help reduce gut inflammation through diverse pathways, including *Bacteroides dorei* (by decreasing gut microbial lipopolysaccharide production^28^), *Faecalibacterium prausnitzii* (through butyrate production^29^) and other *Bifidobacterium* species (*bifidum* and *pseudocatenulatum*, via inhibition of NF-κB activation^30^), and may thus play a role in suppressing *Enterobacteriaceae* growth and CPE colonization^31^.

Pathway analysis based on differential abundance as a function of CPE status providing further supporting evidence that key inflammatory pathways (e.g. sulfate reduction) are enriched during CPE colonization, indicating that they may play a role in the process (**Supplementary Figure 6, Supplementary File 4**). In addition, pathways related to aerobic respiration and oxidative phosphorylation (e.g. pentose phosphate pathway) were also more abundant during CPE colonization consistent with a model of oxygenation of the gut as proposed by Andreas Baumler and Sebastian Winter^32^. In particular, these results were recapitulated after removal of *Enterobacteriaceae* species from functional profiles, highlighting that they are not directly explained by CPE colonization and have substantial contributions from other species as well (**Supplementary Figure 7**). Microaerophilic niches for *Enterobacteriaceae* species due to antibiotic treatment could provide another potential explanation^33^, as antibiotic usage was common in this study (before ∼25% of sampled timepoints, **Supplementary File 1**). As expected, while antibiotic resistance and carbapenemase genes were enriched in gut microbiomes for CPE positive timepoints, no significant differences were observed between index subjects and family members at other timepoints (**Supplementary Figure 8**).

While *Enterobacteriaceae* species were enriched overall in CPE positive samples relative to CPE negative samples (Wilcoxon rank-sum p-value<3.5×10^−5^), index subjects at CPE negative timepoints also showed significantly enriched relative abundances compared to family members (Wilcoxon rank-sum p-value=0.01, **Supplementary Figure 9**). In addition, the composition of *Enterobacteriaceae* species varied across individuals with *Escherichia coli* and *Klebsiella pneumoniae* being the most common species, but other *Escherichia, Klebsiella, Enterobacter* and *Proteus* species also being moderately abundant across some individuals and timepoints (**Supplementary Figure 9**). Of note, while several *Enterobacteriaceae* species exhibited high abundance across individuals, these did not necessarily correspond to the CPE species colonizing a subject (e.g. subject 0505-T in timepoints 1-3). In addition, we observed rapid shifts in *Enterobacteriaceae* profiles (e.g. in 0457-T and 0512-T at timepoint 6) and overall higher variation in *Enterobacteriaceae* abundances across timepoints in index subjects (Wilcoxon rank-sum p-value<0.05; **Figure 1d, Supplementary Figure 9**). Together these results indicate that CPE colonization may be maintained by a altered, dynamic pro-inflammatory microenvironment that supports *Enterobacteriaceae* species, which is resolved in association with recovery of microbiome diversity and function^34^.

### Distinct strain-level dynamics of *Enterobacteriaceae* species in the gut microbiomes of index patients and family members

We next analyzed the deep shotgun metagenomic sequencing data at a higher resolution looking for within-species strain-level dynamics across individuals for the two most prevalent *Enterobacteriaceae* species (*E. coli* and *K. pneumoniae*). Read mapping to reference genomes was used to call high-confidence single-nucleotide variants, and modes in allele frequency distributions were used to infer the number of strains present using a classical approach in population genetics^20,35^ (**Methods, Supplementary Figure 10**). For 53% of the samples (63% for *E. coli*, 38% for *K. pneumoniae*) where a species was confidently detected (relative abundance >0.1%), read coverage was sufficient to identify strain variation (one, two or multiple strains, otherwise classified as low coverage; **Figure 2a, 2b, Supplementary Figure 11**). Overall, as expected for a gut commensal^36,37^, *E. coli* was found at comparable frequencies in index subjects (86%) and family members (90%), and was also more frequently detected in gut microbiome samples overall relative to *K. pneumoniae* (Fisher’s exact p-value<5×10^−20^, **Figure 2a**). *K. pneumoniae* was, however, more frequently found in index subjects (70%) relative to family members (39%), consistent with the hypothesis that a distinct pro-inflammatory environment might be facilitating colonization in these individuals (Fisher’s exact p-value<2×10^−9^, **Figure 2b**).

**Figure 2.**
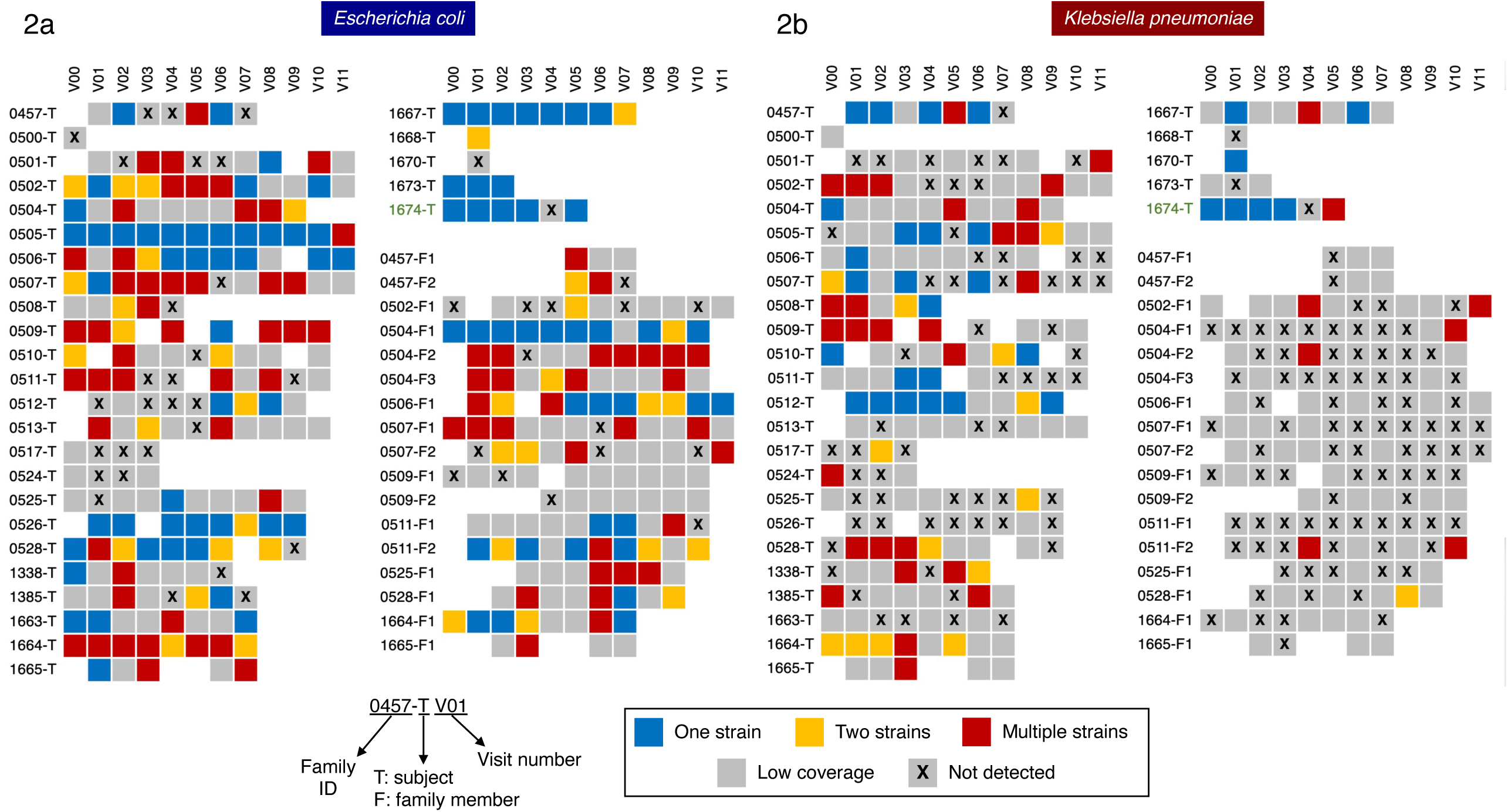
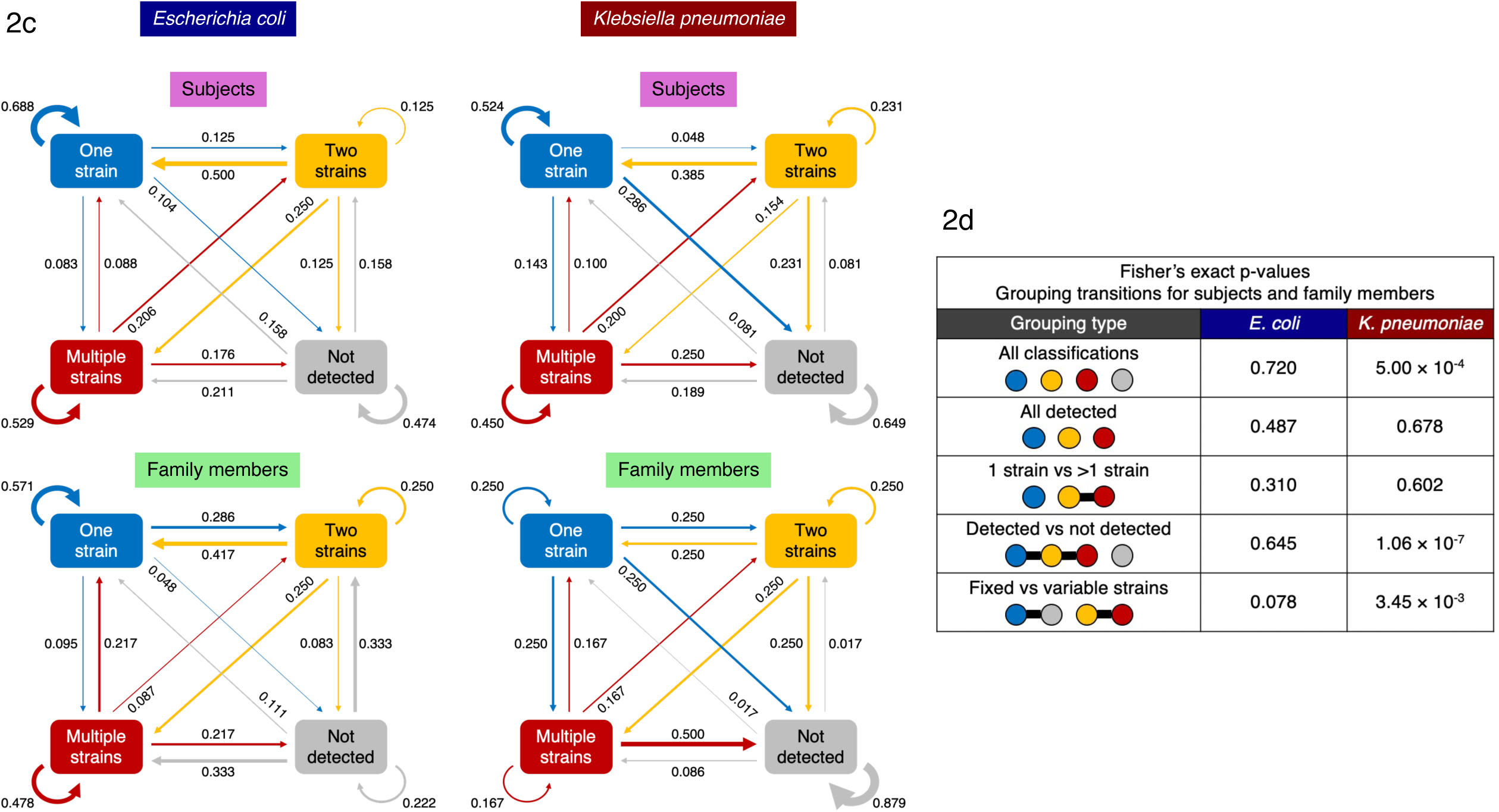
*Enterobacteriaceae* strain variations and dynamics in index subjects and family members. Strain composition for (a) *E. coli* and (b) *K. pneumoniae* determined based on allele frequency distributions per sample. Samples where a species was not detected (relative abundance <0.1%) or in which the genome had low coverage (<10×) were distinguished from these where one, two or more strains where confidently detected. Each row depicts the multiple timepoints for a subject (where available), with index subjects indicated with a T and family members with an F in subject IDs. (c) First-order markov models showing the probability of transition between different states (excluding the low coverage state where information is missing). (d) Table showing statistical significance of differences in transition probabilities between index subjects and family members, for various groupings of states and *Enterobacteriaceae* species. States placed in the same grouping are connected by a horizontal line.

In terms of strain variations, of the samples that were assigned a classification, we noted that *E. coli* was frequently observed as a single distinct strain in the gut microbiome of index patients (44%) while family members more often had multiple strains (44%; **Supplementary Figure 11**), suggesting that a single clone may often dominate in a pro-inflammatory environment. A few individuals also maintained a single strain state over the course of several months (up to a year, e.g. 0505-T, 1667-T, 0506-T) indicating that this can be a stable state for some individuals (**Figure 2a**). For *K. pneumoniae*, despite being detected more sporadically in index subjects and family members, the multi-strain state was the more common observation (49%), consistent with the hypothesis that even in a pro-inflammatory environment no distinct clone will typically outcompete others^38^ (**Figure 2b, Supplementary Figure 11**). Overall, in agreement with our previous observations (**Figure 1d, Supplementary Figure 9**), we noted that strain compositions were highly variable for these *Enterobacteriaceae* species over time.

Capturing transition frequencies between various strain compositions as a first-order Markov model (maximum likelihood with Laplace smoothing), we noted distinct patterns for *E. coli* and *K. pneumoniae*, as well as between index subjects and family members (**Figure 2c, Methods**). For example, *E. coli* colonization is more likely to stay in a single strain state for index patients (69%), relative to family members (57%), as well as relative to single strain *K. pneumoniae* colonization (52%, **Figure 2c**). Also, when *E. coli* is not detected, this state is more likely to be maintained in index subjects (47%) than in family members (22%, **Figure 2c**). Overall the Markov model predicts that *E. coli* in index subjects and family members tend to be in the one strain state (41% for subjects, 39% for family members). In contrast, *K. pneumoniae* frequently converges to the not detected state in subjects (42%) and in family members (75%). Grouping various classes of detection and strain status in different ways, we then tested if index subjects and family members show different transition probabilities in *E. coli* or *K. pneumoniae* (**Figure 2d**). For *E. coli*, transition probabilities were not significantly different between index subjects and family members (Fisher’s exact p-value>0.05, **Figure 2d**). In contrast, driven by the stark detected/not detected patterns seen for *K. pneumoniae*, index subjects had significantly different transition probabilities compared to family members for various groupings that involve the not detected state (“All classifications”, “Detected vs not detected” and “Fixed vs variable strains”, Fisher’s exact p-value<10^−2^, **Figure 2d**). These results further highlight the differences in strain-level dynamics for *Enterobacteriaceae* species in the potentially pro-inflammatory gut microbiome milieu of index subjects.

### Sub-strain variation and plasmid sharing in *Enterobacteriaceae* species in relation to CPE decolonization

Samples that were determined to have a single-strain can nevertheless exhibit sub-strain variation in relation to this genomic background, similar to quasi-species diversity in viral populations. Characterizing the distribution of such intra-host variations across genes can help identify adaptive changes that may be important for CPE colonization, similar to recent studies with mouse models and strain isolates^39,40^. To analyze this standing variation in *Enterobacteriaceae* species, we identified low-frequency (<50%) single-nucleotide variants in single-strain timepoints (30,155 and 13,061 SNVs for *E. coli* and *K. pneumoniae*, respectively), and analyzed them for protein function altering changes to identify potential adaptive changes in the genomes of *Enterobacteriaceae* strains during gut colonization (**Supplementary File 5, Methods**). In total we found 5,919 and 1,787 putative function altering changes in *E. coli* and *K. pneumoniae*, including several in key polysaccharide utilization and virulence (e.g. *lacZ, lacY*, ECIAI39_4258 [Putative invasin/intimin protein]) similar to what has been described based on isolate sequencing as being key genes undergoing selection for colonization of the human gut^41,42^ (**Table 2**). In particular, we visualized function-altering SNVs in genes implicated in polysaccharide utilization, where adaptive mutations can reflect pressures to make use of polysaccharides derived from the host diet, to identify several structural motifs that might be key to their function (**Supplementary Figure 12**). Consistent with the fact that we are studying low-frequency SNVs, we noted that most regions bear signatures of purifying selection for these SNVs (dN/dS<0.5, **Supplementary Figure 12**), though overall the identified genes were significantly enriched relative to the genome-wide average for non-synonymous SNVs (**Table 2**).

To study these variations further in relation to CPE decolonization, the time-series information was used to cluster SNVs that co-vary (**Methods**). Interestingly, in some subjects multiple clusters were revealed by this analysis, indicating that there were distinct sub-strain lineages that differed by a few hundred SNVs genome-wide (e.g. 1674-T, **Figure 3a, b**; **Supplementary Figure 13-15**). In particular for subject 1674-T, we noted that both *E. coli* and *K. pneumoniae* have a dominant cluster during CPE positive timepoints (V00–V03) that match the SNV signature seen in the genomes for *E. coli* and *K. pneumoniae* CPE isolates for this individual (*Shared* and *Cluster 1* SNVs, **Figure 3c, d, Supplementary Figure 15**). In contrast, the sub-dominant cluster (*Cluster 2*, **Figure 3c, d, Supplementary Figure 15**; likely representing a sub-lineage of *Cluster 1*) has a SNV signature that is not seen in the CPE isolates and is still detected in the post-decolonization timepoint (V05, based on stool PCR testing), indicating that these sub-strain lineages may be discordant for CPE status despite their overall genomic similarity. For both *E. coli* and *K. pneumoniae*, we noted that decolonization coincides with the appearance of a distinct strain with >1,000 SNVs distinguishing them from the CPE strains (*V05 unique*, **Figure 3c, d, Supplementary Figure 15**; V05 classified as two-strain timepoint). Interestingly, despite these shared patterns within *E. coli* and *K. pneumoniae* strains, we noted that they exhibited dissimilar trends in terms of overall relative abundance, with the abundance of *E. coli* being reduced leading up to the decolonization timepoint (V05) while *K. pneumoniae* abundance peaks at this point (**Figure 3e, f**). In general, while a few dense trajectories of co-varying SNVs were detected in other individuals, many SNVs varied independent of these clusters (**Supplementary Figure 13, 14**). Overall, these results suggest that *Enterobacteriaceae* species may share patterns of sub-strain dynamics in relation to CPE decolonization, despite having species-specific ecological properties.

**Figure 3.**
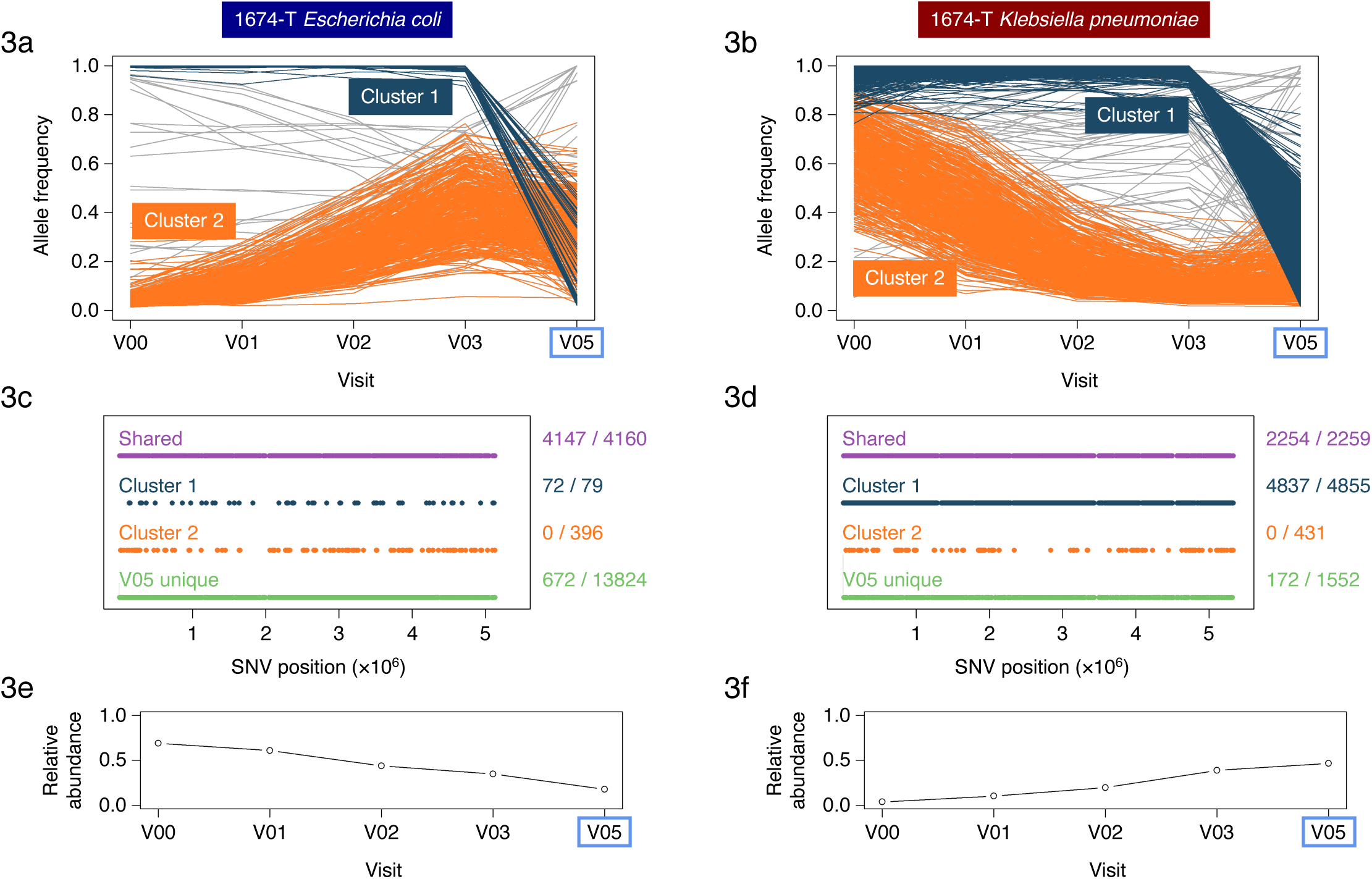
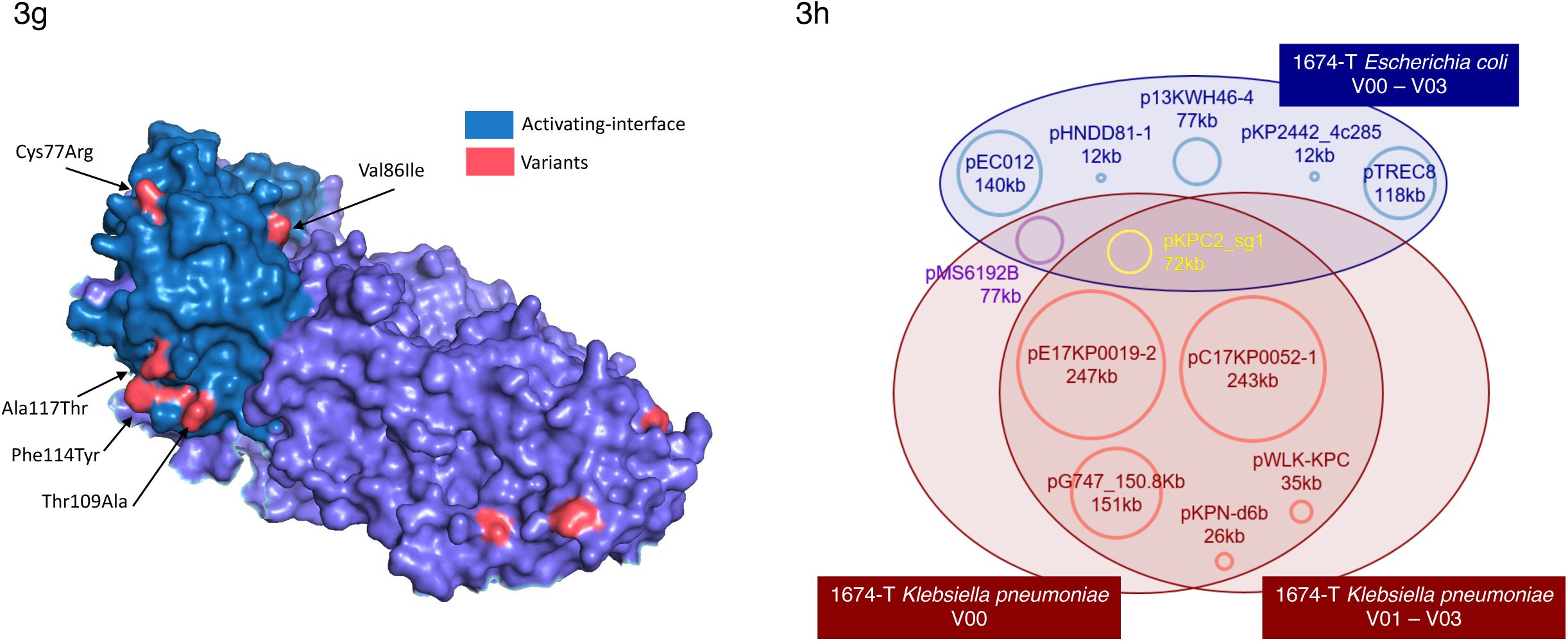
Sub-strain variation and plasmid sharing in *Enterobacteriaceae* species. Line charts showing the allele frequencies of metagenomics-derived SNVs across timepoints for (a) *E. coli* and (b) *K. pneumoniae*, for subject 1674-T. SNVs that belong to different sub-strain clusters are colored accordingly and a potential model for the corresponding haplotypes is shown in **Supplementary Figure 15**. All genome-wide SNVs detected at all the depicted timepoints are shown here. The blue box indicates the CPE negative timepoint. (c, d) Corresponding genomic distribution for SNVs belonging to cluster 1, cluster 2, and those that are fixed at all timepoints (“Shared”) or just in the CPE negative timepoint (“V05 unique”). The associated fraction describes the number of SNVs in a grouping that are common with CPE isolates, in relation to the total number of SNVs in this grouping. (e, f) Relative abundance of *E. coli* and *K. pneumoniae* in the gut metagenome for subject 1674-T across timepoints. (g) Visualization of amino acid changes (red) seen in a key polysaccharide utilization protein (lacZ, pdb structure 4DUX chain A, identified based on **Table 2, 3**) based on metagenomics-derived SNVs that changed in frequency over time and across multiple individuals. SNVs are found primarily on the protein surface, with six SNVs located on the activating interface responsible for tetramerization (teal). (h) Diagram depicting plasmid sharing between *E. coli* and *K. pneumonia*e strains at various time points for subject 1674-T. Plasmid sequences were clustered at 95% similarity (**Supplementary Figure 16**) and a representative plasmid for each cluster is shown in the Venn diagram as an approximately sized circle, plasmid name and size.

Leveraging the availability of multiple timepoints across subjects, we identified SNVs whose populations frequencies varied notably over time (>30%). These were then overlapped across subjects to identify SNVs that have this property recurrently (**Table 3, Supplementary File 6**), identifying a range of polysaccharide utilization (*lacZ, lacI, treA*), pyruvate metabolism (*pflB, pykF*) and protein synthesis (*dnaK*, 30S and 50S ribosomal subunits) genes that have been implicated in adaptive evolution under nutrient limitation^43^, antibiotic^44^ and environmental stress^45,46^ conditions. In particular, several genes were common to the lists for *E. coli* and *K. pneumoniae* (*srmB, pnp, nlpI, pheT*), suggesting that similar selection constraints might be acting on strains for both species. The *lacZ* gene was highlighted as having the most recurrent, frequency-varying SNVs in this analysis (n=12), with all SNVs occurring in surface-exposed regions (**Figure 3g**). Comparing the accessible surface area (ASA) of protein residues between variant and other sites revealed that variant residues are significantly more exposed to the solvent (mean=73.1Å^2^) than other residues (mean=34.1Å^2^, Welch’s t-test p-value <0.01). In addition, 6 SNVs occurred in the activating interface, a region near the amino-terminus of lacZ that is required for tetramerization^47^, indicating that they may influence lacZ function via complex formation dynamics.

Among the genetic features prominent in CPE strains seen in subject 1674-T, we noted that variants in polysaccharide utilization genes were common as discussed previously (**Figure 3g**). In addition, we analyzed plasmid sequences across timepoints and identified two important plasmids that were shared between *E. coli* and *K. pneumoniae* CPE strains (**Figure 3h, Supplementary Figure 16, Methods**). This included the pKPC2 plasmid that was recently identified in hypervirulent, carbapenem-resistant *Klebsiella pneumoniae* isolates from Singapore and harbors *bla*_KPC-2_, a carbapenemase gene that was the basis of CPE designation for these isolates^24^. In addition, the pMS6192B plasmid was shared between all *E. coli* isolates and the *Klebsiella pneumoniae* isolate from the first visit (V00, **Figure 3h**). The shared plasmids have a total sequence length of >140kbp and no SNVs distinguishing the two species, indicating that they have a recent common source. Plasmid transfer experiments with pKPC2 between *E. coli* and *K. pneumoniae* strains suggest moderate conjugation frequency under *in vitro* conditions (∼0.1%, **Methods**). In addition, half of the plasmid bearing clones (3/6) were observed to have a SNV in pKPC2 after 300 generations, defining an upper-bound on the divergence of plasmid-bearing isolates having no SNVs being 5 months (Binomial p-value <0.05).

## Discussion

The availability of metagenomic data from up to 12 timepoints over the period of a year in this longitudinal study allowed us examine long-term dynamics, enabling comparison of microbiome configurations before and after CPE decolonization in a subject-matched fashion to reveal microbiome shifts associated with decolonization. This analysis revealed ecological shifts that cannot be explained solely by the loss of CPE strains (e.g. increase in species richness), and the specific taxonomic and functional changes observed point to the role of inflammation in maintaining an *Enterobacteriaceae*-favorable gut environment in index subjects (e.g. *Pantoea* species; **Supplementary Figure 2, Supplementary File 3**). In addition, our data indicates that this configuration may be unstable in many individuals, opening up the possibility that interventions that reduce gut inflammation directly or via the action of probiotics could reduce *Enterobacteriaceae* abundances and promote CPE decolonization.

In particular, gut inflammation has been known to create a niche for enterics such as *Salmonella*^48^, where some species can use sulfate, nitrate and tetrathionate as the terminal electron acceptor for anaerobic respiration (e.g. *E. coli*). The enriched pathways in CPE colonized subjects are marked by menaquinol biosynthesis, glycolysis and respiration (TCA cycle), even after computationally subtracting out the contribution of *Enterobacteriaceae*, indicating that the gut environment in this group is qualitatively different in oxygenation. In addition, fucose and rhamnose degradation, as well as 1, 2-propanediol degradation are enriched in CPE colonization, potentially serving as carbon sources for *Enterobacteriaceae* such as *K. pneumoniae* which can demonstrate competitive fitness in the gut with oxygen as terminal electron acceptor under such conditions^38^. The enrichment of the pentose phosphate pathway could indicate a need for reducing equivalents of NADPH^+^ to maintain redox conditions or serve as nucleic acid precursors to fuel growth. Overall, the shift in microbial pathways in CPE colonized subjects appears to be largely independent of *Enterobacteriaceae* species, but favoring their growth. Further work is needed to understand if this shift is primarily established by gut inflammation (e.g. as seen in colitis^49^, potentially through direct measurement of protein biomarkers such as Calprotectin) or if a diverse set of factors play a strong role in an individual-specific manner (e.g. antibiotics for some subjects^33^). In particular, while the reduction in microbial diversity during CPE colonization could not be solely attributed in this study to factors such as antibiotic usage or hospitalization at a timepoint, these could be delayed effects and would therefore need a more controlled study design to explore this further.

An alternative strategy to promote CPE decolonization could be based on the introduction of species that were relatively depleted in the colonized state (e.g. *Faecalibacterium prausnitzii* or *Bifidobacterium bifidum*), either in the form of probiotic formulations or through the use of fecal microbiota transplants^50^. Matching donors to recipients to supplement missing species or to promote further instability in *Enterobacteriaceae* abundances based on ecological models could be a promising avenue to explore here similar to studies for *Clostridoides difficile*^51,52^. The observed differences in colonization dynamics for *Enterobacteriaceae* species (*E. coli* and *K. pneumoniae*) suggest that CPE decolonization strategies might also have to be species-specific. For example, the presence of multiple *K. pneumoniae* strains in index subjects is consistent with the hypothesis that they are not well-adapted for gut colonization but are instead opportunistically exploiting a niche. Decolonization of *K. pneumoniae* strains may therefore require elimination of conditions that favor this niche such as inflammation or availability of specific sugars. On the other hand, the presence of a single strain of *E. coli* in many index subjects supports a model where gut adapted strains have acquired antimicrobial resistance cassettes, and plasmid targeting strategies might be better suited in this case. Interestingly, data from our cohorts suggests that human gut microbiomes can harbor multiple strains of commensal species such as *E. coli* (in contrast to observations in mouse studies^53,54^, even among non-CPE colonized family members^55,56^ (**Figure 2a**). Further studies using high-throughput culturing and single-cell sequencing could help accurately reconstruct strain genomes and unravel the factors that determine niche competition^57^.

Understanding the factors that support gut colonization by CPE species can provide another avenue to identify targets for intervention. As we show here, the analysis of high-coverage metagenomic data to identify sub-strain variations with functional impact can provide promising hypotheses based on *in vivo* evolution, similar to the quasi-species analysis of viruses^58,59^, or mutagenesis-based experiments^60^. Furthermore, identification and isolation of sub-strain lineages with distinct advantages in colonizing the host or avoiding decolonization (e.g. as may be the case for cluster 2 in 1674-T), can help narrow down the genetic features that need to be investigated *in vitro*. Finally, the role of the gut microbiome as a reservoir for AMR determinants, and plasmid sharing across *Enterobacteriaceae* species is of particular concern. While we cannot definitively conclude that the data for index subject 1674-T represents an example of plasmid transfer, these observations and the isolated strains serve as important resources to guide further investigations into plasmid transmission and CPE decolonization.

## Methods

### Sample collection and CPE classification

A prospective cohort study involving CPE carriers was conducted from October 2016 to February 2018. Study participants were recruited from two tertiary healthcare centers in Singapore. CPE carriers were identified by routine collection of rectal swab samples for clinical care and infection prevention and control measures, in accordance with local infection control policies. The study received ethics approval from the Singapore National Healthcare Group Domain Specific Review Board 74 (NHG DSRB Reference: 2016/00364) prior to commencement. Stool samples were first collected weekly for four weeks, then monthly for five months, and finally once every two months for six months. In addition to the CPE-colonized subjects, stool samples from a number of family members were also obtained to provide a control dataset. Samples obtained from index subjects were classified as either CPE positive or CPE negative, based on whether CPE genes (including *bla*_NDM-1_, *bla*_KPC_, *bla*_OXA-48_, *bla*_IMI-1_, and *bla*_IMP_) were positively identified from *Enterobacteriaceae* isolates found to be resistant to either meropenem or ertapenem^24^ (**Supplementary Table 1**). The presence of CPE negative samples was used to detect CPE clearance and samples were further classified based on the amount of time elapsed since clearance i.e. before clearance, within two months post-clearance, and more than two months post-clearance. Due to the focus on household transmission and CPE carriage, dietary information was not collected in the clinical study.

### Isolate sequencing and assembly

DNA for all CPE isolates obtained from stool samples in this study (all subjects, all timepoints) was collected from Tan Tock Seng Hospital (TTSH) and transferred to the Genome Institute of Singapore (GIS) for whole genome sequencing. Library preparation was performed using the NEBNext Ultra DNA Library Prep Kit for Illumina, and 2×151 base-pair sequencing was performed using the Illumina HiSeq 4000. Raw FASTQ reads were processed using in-house pipelines at GIS for *de novo* assembly with the Velvet assembler^61^ (v1.2.10), parameters optimized by Velvet Optimiser (*k*-mer length ranging from 81 to 127), contig scaffolding with Opera^62^ (v1.4.1), and finishing with FinIS^63^ (v0.3).

### Shotgun metagenomic sequencing

DNA from 361 stool samples was extracted using the PowerSoil DNA Isolation Kit (12888, MoBio Laboratories) with modifications to the manufacturer’s protocol. Specifically, to avoid spin filter clogging, we extended the centrifugation to twice the original duration, and solutions C2, C3 and C4 were doubled in volume. DNA was eluted in 80µL of Solution C6. Concentration of DNA was determined by Qubit dsDNA BR assay (Q32853, Thermo Fisher Scientific). For library construction, 50ng of DNA was re-suspended in a total volume of 50µL, and was sheared using Adaptive Focused Acoustics (Covaris) with the following parameters: duty factor of 30%, peak incident power (PIP) of 450, 200 cycles per burst, and treatment time of 240s. Sheared DNA was cleaned up with 1.5× Agencourt AMPure XP beads (A63882, Beckman Coulter). Gene Read DNA Library I Core Kit (180434, Qiagen) was used for end-repair, A-addition and adapter ligation. Custom barcode adapters were used for cost considerations (HPLC purified, double stranded, 1^st^ strand: 5’ P-GATCGGAAGAGCACACGTCT; 2^nd^ strand: 5’ ACACTCTTTCCCTACACGACGCTCTTCCGATCT) in replacement of Gene Read Adapter I Set for library preparation. Library was cleaned up twice using 1.5× Agencourt AMPure XP beads (A63882, Beckman Coulter). Enrichment was carried out with indexed-primers according to an adapted protocol from Multiplexing Sample Preparation Oligonucleotide kit (Illumina). We polled the enriched libraries in equi-molarity and sequenced them on an Illumina HiSeq 2500 sequencing instrument at GIS to generate 2×101 base-pair reads, which yielded around 17.7 billion paired-end reads in total and 49 million paired-end reads on average per library.

### Taxonomic and functional profiling

Read quality trimming was performed using famas (https://github.com/andreas-wilm/famas, v0.10, --no-order-check), and microbial reads were identified by mapping and filtering out reads aligned to the human reference genome (hg19) using bwa-mem^64^ (v0.7.9a, default parameters; >90% microbial reads on average). Taxonomic profiling was done using MetaPhlAn^65^ (v2.0, default parameters, filtering taxa with relative abundance<0.1%) and functional profiles were obtained with HUMAnN^66^ (v2.0, default parameters). As a sanity check, we confirmed that species and genus-level taxonomic profiles were not dominated by taxa that are commonly attributed to reagent or laboratory contamination^67^ (**Supplementary File 2**). Average-linkage hierarchical clustering of taxonomic profiles was used to group samples with the number of clusters determined using Akaike information criterion (AIC). Sample α-diversity was computed using the Shannon diversity index with the vegan library in R. Differential abundance analysis was performed using LEfSe^68^ (v1.0.8), as a non-parametric and conservative approach to identify significantly varying taxa and functions across groups^69^. These results were further validated using Songbird^70^ (v1.0.3; –epochs 10000 –differential-prior 0.5) analysis with Bonferroni-corrected p-value<0.05. Abundances of antibiotic resistance genes (ARGs) in the metagenomes was computed using a direct read mapping approach implemented in SRST2^71^ with default parameters and the CARD_v3.0.8_SRST2 database^72^.

### Linear mixed-effects modeling

Linear mixed effects modeling was conducted using the *lmer* function from the *lme4* package in R. For each model, genus-level Shannon diversity was set as the response variable, with colonization status as the fixed effect and potential confounders (e.g. antibiotic usage since last visit, hospitalization status, individual subjects, gender and ethnicity; **Supplementary File 1**) as random effect covariates. Residual Shannon diversity values were derived for visualization by subtracting the intercept terms corresponding to random effects.

### Single-nucleotide variant analysis

Genome assemblies were aligned to their respective reference genomes using nucmer (v3.23, -maxmatch -nosimplify) and consensus SNVs were called using the show-SNVs function in MUMmer^73^. References for *E. coli* (NC_011750) and *K. pneumoniae* (NC_016845.1) were selected to minimize median distance from isolate genomes. Metagenomic SNVs (consensus and low-frequency) were identified based on read mapping using bwa-mem^64^ to the *E. coli* and *K. pneumoniae* references (v0.7.10a; soft-clipped reads and reads with >3 or 4 mismatches for *K pneumoniae* and *E. coli* respectively were filtered out to avoid mis-mapped reads) and variant calling with LoFreq^74^ (v1.2.1; default parameters). Note that our stringent mapping approach restricts to only reads with >96% identity with the reference, and thus will typically exclude mis-mapping of reads from other genomes. Additionally, genomic regions with frequent ambiguous mappings were identified based on isolate sequencing data and metagenome data from samples without target species as determined from taxonomic classification (>5× coverage with *E. coli* reads on *K. pneumoniae* genome or vice versa). Calls in these regions that match positions where variants were called between isolate reads and reference sequence (allele frequency > 95%) were excluded from downstream analysis. The validity of this pipeline was confirmed by noting that very few *K. pneumoniae* SNVs (median=2, mean=5.5) were called genome-wide when analysing metagenomes where taxonomic profiling detected few *K. pneumoniae* reads (10 samples with 107-288 reads). Note that SNVs from such “low coverage” samples are also excluded from further analysis in this study as defined below. To assess the impact of a shared, but potentially divergent, reference on SNV calling, reads were also mapped onto CPE isolate genomes (where available) to call SNVs and compute concordance. Isolate genome based SNVs were translated to the common reference coordinate system using the UCSC liftover tool^75^ with chain file generated using flo^76^ (-fastMap -tileSize=12 -minIdentity=90).

### Strain analysis

Metagenomic coverage of samples for *E. coli* and *K. pneumoniae* was determined from bwa-mem read mappings using genomeCoverageBed^77^ (v2.25.0). Samples with too low relative abundance for confident identification (<0.1%) were designated as “not detected”, while samples with low median read coverage (<8) were designated as “low coverage”. Of the remaining samples, those with >90% of the SNVs at or above an allele frequency of 0.9 were designated as “one strain”, exhibiting a unimodal distribution as is classically expected in the single haplotype setting^20^ (**Supplementary Figure 10a**). A *k*-means clustering approach (based on allele frequency values < 0.98, *k*=2) was used on other samples to identify “two strains” (silhouette score > 0.8, indicating good concordance with 2 clusters for a bimodal distribution) and “multiple strains” cases where there may be more than 2 clusters (**Supplementary Figure 10a**). Note that this analysis was only used to determine strain “states” (**Figure 2**), and the corresponding clusters were not used for downstream haplotype analysis. To confirm metagenomic SNV calling quality and strain designations, “one strain” cases were compared to SNVs from corresponding isolates (where available) and noted to have high precision for both *E. coli* and *K. pneumoniae* (>98%, **Supplementary Figure 10b**). A first-order Markov model of the transition frequencies between the strain compositions was estimated using the markovchain package in R^78^ (maximum likelihood estimator with Laplace smoothing parameter = 1).

### Sub-strain analysis

SNVs with mean allele frequency >0.9 across timepoints were identified as likely fixed across all strains in a sample. Non-fixed SNVs from “one strain” cases were further annotated for their impact on protein function using SnpEff^79^ (v4.3). The ratio of the rate of non-synonymous (dN) to synonymous (dS) mutations was calculated using the package Biopython.codonalign.codonseq with the ‘NG86’ method.

Leveraging the availability of multiple “one strain” timepoints in some individuals, non-fixed SNV trajectories were clustered to identify co-varying SNVs that may belong to a common sub-strain background. Specifically, the DBSCAN algorithm in R^80^ was used to cluster SNV trajectories in selected individuals with multiple “one strain” timepoints (ε=0.2, minPts=2*n* as recommended) and identified clusters were visualized as a sanity check.

### Plasmid analysis

A Mash screen search approach was used with PLSDB^81^ to obtain a list of plasmids that are potentially present in the CPE isolate genomes. The union of all such plasmid sequences was then aligned with isolate genome assemblies to identify plasmids hits with >85% coverage at 95% identity (only alignments >500bp). Plasmid hits were clustered into groups using hierarchical clustering at 95% identity (hclust function in R, average linkage based on Mash distance^82^), with the longest plasmid serving as a representative. Only plasmids longer than 10kbp are included in the figure to avoid spurious/redundant matches to shorter plasmids.

### Plasmid conjugation assay

Donor *E. coli* harbouring the pKPC2 plasmid with a kanamycin selection cassette (MG1655) and recipient *K. pneumoniae* strains (ATCC13883) were streaked on selective LBA and incubated overnight at 37°C. Bacterial colonies were resuspended in LB (1 mL), diluted to OD_600_ = 0.5 and mixed in a 1:1 ratio and spotted onto 0.22 µm nitrocellulose membrane (Sartorius) placed on top of LBA (20 µL). After 4 hours incubation at 37°C, the bacterial mixture was resuspended in 2 mL of PBS, serially diluted and plated on LBA with appropriate antibiotic selection. Kanamycin (50 micrograms/ml) and fosfomycin (40 micrograms/ml) were used for selection of transconjugants. Plates were incubated at 37°C overnight and colonies were enumerated. Conjugation frequency was calculated as the total number of transconjugants per total number of recipients.

### Code and data availability

Source code for scripts used to analyze the data are available in a GitHub project at https://github.com/CSB5/CPE-microbiome. Isolate and shotgun metagenomic sequencing data is available from the European Nucleotide Archive (ENA – https://www.ebi.ac.uk/ena/browser/home) under project accession number PRJEB49334.

## References

1. von Wintersdorff CJ, Penders J, van Niekerk JM, et al. Dissemination of Antimicrobial Resistance in Microbial Ecosystems through Horizontal Gene Transfer. Front Microbiol. 2016;7:173. doi:10.3389/fmicb.2016.00173

2. van Duin D, Doi Y. The global epidemiology of carbapenemase-producing Enterobacteriaceae. Virulence. 05 2017;8(4):460–469. doi:10.1080/21505594.2016.1222343

3. Suay-García B, Pérez-Gracia MT. Present and future of carbapenem-resistant Enterobacteriaceae (CRE) infections. Antibiotics (Basel). Aug 2019;8(3):122. doi:10.3390/antibiotics8030122

4. Blair JM, Webber MA, Baylay AJ, Ogbolu DO, Piddock LJ. Molecular mechanisms of antibiotic resistance. Nat Rev Microbiol. Jan 2015;13(1):42–51. doi:10.1038/nrmicro3380

5. Exner M, Bhattacharya S, Christiansen B, et al. Antibiotic resistance: What is so special about multidrug-resistant Gram-negative bacteria? GMS Hyg Infect Control. 2017;12:Doc05. doi:10.3205/dgkh000290

6. Papp-Wallace KM, Endimiani A, Taracila MA, Bonomo RA. Carbapenems: past, present, and future. Antimicrob Agents Chemother. Nov 2011;55(11):4943–60. doi:10.1128/AAC.00296-11

7. Codjoe FS, Donkor ES. Carbapenem Resistance: A Review. Med Sci (Basel). Dec 2017;6(1)doi:10.3390/medsci6010001

8. Calfee D, Jenkins SG. Use of active surveillance cultures to detect asymptomatic colonization with carbapenem-resistant Klebsiella pneumoniae in intensive care unit patients. Infect Control Hosp Epidemiol. Oct 2008;29(10):966–8. doi:10.1086/590661

9. Schechner V, Kotlovsky T, Kazma M, et al. Asymptomatic rectal carriage of blaKPC producing carbapenem-resistant Enterobacteriaceae: who is prone to become clinically infected? Clin Microbiol Infect. May 2013;19(5):451–6. doi:10.1111/j.1469-0691.2012.03888.x

10. Penders J, Stobberingh EE, Savelkoul PH, Wolffs PF. The human microbiome as a reservoir of antimicrobial resistance. Front Microbiol. 2013;4:87. doi:10.3389/fmicb.2013.00087

11. Nordmann P, Naas T, Poirel L. Global spread of Carbapenemase-producing Enterobacteriaceae. Emerg Infect Dis. Oct 2011;17(10):1791–8. doi:10.3201/eid1710.110655

12. Haidar G, Clancy CJ, Chen L, et al. Identifying Spectra of Activity and Therapeutic Niches for Ceftazidime-Avibactam and Imipenem-Relebactam against Carbapenem-Resistant Enterobacteriaceae. Antimicrob Agents Chemother. 09 2017;61(9)doi:10.1128/AAC.00642-17

13. Tooke CL, Hinchliffe P, Bragginton EC, et al. β-Lactamases and β-Lactamase Inhibitors in the 21st Century. J Mol Biol. 08 2019;431(18):3472–3500. doi:10.1016/j.jmb.2019.04.002

14. Sun X, Winglee K, Gharaibeh RZ, et al. Microbiota-Derived Metabolic Factors Reduce Campylobacteriosis in Mice. Gastroenterology. 05 2018;154(6):1751–1763.e2. doi:10.1053/j.gastro.2018.01.042

15. Ichinohe T, Pang IK, Kumamoto Y, et al. Microbiota regulates immune defense against respiratory tract influenza A virus infection. Proc Natl Acad Sci U S A. Mar 2011;108(13):5354–9. doi:10.1073/pnas.1019378108

16. Buffie CG, Bucci V, Stein RR, et al. Precision microbiome reconstitution restores bile acid mediated resistance to Clostridium difficile. Nature. Jan 2015;517(7533):205–8. doi:10.1038/nature13828

17. Haak BW, Littmann ER, Chaubard JL, et al. Impact of gut colonization with butyrate-producing microbiota on respiratory viral infection following allo-HCT. Blood. 06 2018;131(26):2978–2986. doi:10.1182/blood-2018-01-828996

18. Lieberman TD, Michel JB, Aingaran M, et al. Parallel bacterial evolution within multiple patients identifies candidate pathogenicity genes. Nat Genet. Nov 2011;43(12):1275–80. doi:10.1038/ng.997

19. Lieberman TD, Flett KB, Yelin I, et al. Genetic variation of a bacterial pathogen within individuals with cystic fibrosis provides a record of selective pressures. Nat Genet. Jan 2014;46(1):82–7. doi:10.1038/ng.2848

20. Garud NR, Good BH, Hallatschek O, Pollard KS. Evolutionary dynamics of bacteria in the gut microbiome within and across hosts. PLoS Biol. 01 2019;17(1):e3000102. doi:10.1371/journal.pbio.3000102

21. Davenport ER, Sanders JG, Song SJ, Amato KR, Clark AG, Knight R. The human microbiome in evolution. BMC Biol. 12 2017;15(1):127. doi:10.1186/s12915-017-0454-7

22. Chu ND, Smith MB, Perrotta AR, Kassam Z, Alm EJ. Profiling Living Bacteria Informs Preparation of Fecal Microbiota Transplantations. PLoS One. 2017;12(1):e0170922. doi:10.1371/journal.pone.0170922

23. Ferreiro A, Crook N, Gasparrini AJ, Dantas G. Multiscale Evolutionary Dynamics of Host-Associated Microbiomes. Cell. 03 2018;172(6):1216–1227. doi:10.1016/j.cell.2018.02.015

24. Mo Y, Hernandez-Koutoucheva A, Musicha P, et al. Duration of Carbapenemase-Producing Enterobacteriaceae Carriage in Hospital Patients. Emerg Infect Dis. Sep 2020;26(9):2182–2185. doi:10.3201/eid2609.190592

25. Haverkate MR, Weiner S, Lolans K, et al. Duration of Colonization With Klebsiella pneumoniae Carbapenemase-Producing Bacteria at Long-Term Acute Care Hospitals in Chicago, Illinois. Open Forum Infect Dis. Oct 2016;3(4):ofw178. doi:10.1093/ofid/ofw178

26. Lewis JD, Enfield KB, Mathers AJ, Giannetta ET, Sifri CD. The limits of serial surveillance cultures in predicting clearance of colonization with carbapenemase-producing Enterobacteriaceae. Infect Control Hosp Epidemiol. Jul 2015;36(7):835–7. doi:10.1017/ice.2015.57

27. Korach-Rechtman H, Hreish M, Fried C, et al. Intestinal Dysbiosis in Carriers of Carbapenem-Resistant. mSphere. 04 2020;5(2)doi:10.1128/mSphere.00173-20

28. Yoshida N, Emoto T, Yamashita T, et al. Bacteroides vulgatus and Bacteroides dorei Reduce Gut Microbial Lipopolysaccharide Production and Inhibit Atherosclerosis. Circulation. 11 2018;138(22):2486–2498. doi:10.1161/CIRCULATIONAHA.118.033714

29. Lenoir M, Martín R, Torres-Maravilla E, et al. Butyrate mediates anti-inflammatory effects of. Gut Microbes. Nov 2020;12(1):1–16. doi:10.1080/19490976.2020.1826748

30. Riedel CU, Foata F, Philippe D, Adolfsson O, Eikmanns BJ, Blum S. Anti-inflammatory effects of bifidobacteria by inhibition of LPS-induced NF-kappaB activation. World J Gastroenterol. Jun 2006;12(23):3729–35. doi:10.3748/wjg.v12.i23.3729

31. Zeng MY, Inohara N, Nuñez G. Mechanisms of inflammation-driven bacterial dysbiosis in the gut. Mucosal Immunol. 01 2017;10(1):18–26. doi:10.1038/mi.2016.75

32. Winter SE, Bäumler AJ. A breathtaking feat: to compete with the gut microbiota, Salmonella drives its host to provide a respiratory electron acceptor. Gut Microbes. 2011 Jan-Feb 2011;2(1):58–60. doi:10.4161/gmic.2.1.14911

33. Rivera-Chávez F, Lopez CA, Bäumler AJ. Oxygen as a driver of gut dysbiosis. Free Radic Biol Med. 04 2017;105:93–101. doi:10.1016/j.freeradbiomed.2016.09.022

34. Chng KR, Ghosh TS, Tan YH, et al. Metagenome-wide association analysis identifies microbial determinants of post-antibiotic ecological recovery in the gut. Nat Ecol Evol. 09 2020;4(9):1256–1267. doi:10.1038/s41559-020-1236-0

35. Anyansi C, Straub TJ, Manson AL, Earl AM, Abeel T. Computational Methods for Strain-Level Microbial Detection in Colony and Metagenome Sequencing Data. Front Microbiol. 2020;11:1925. doi:10.3389/fmicb.2020.01925

36. Tenaillon O, Skurnik D, Picard B, Denamur E. The population genetics of commensal Escherichia coli. Nat Rev Microbiol. Mar 2010;8(3):207–17. doi:10.1038/nrmicro2298

37. Bailey JK, Pinyon JL, Anantham S, Hall RM. Commensal Escherichia coli of healthy humans: a reservoir for antibiotic-resistance determinants. J Med Microbiol. Nov 2010;59(Pt 11):1331–1339. doi:10.1099/jmm.0.022475-0

38. Stacy A, Andrade-Oliveira V, McCulloch JA, et al. Infection trains the host for microbiota-enhanced resistance to pathogens. Cell. Feb 2021;184(3):615–627.e17. doi:10.1016/j.cell.2020.12.011

39. Barreto HC, Sousa A, Gordo I. The Landscape of Adaptive Evolution of a Gut Commensal Bacteria in Aging Mice. Curr Biol. 03 23 2020;30(6):1102–1109.e5. doi:10.1016/j.cub.2020.01.037

40. Ernst CM, Braxton JR, Rodriguez-Osorio CA, et al. Adaptive evolution of virulence and persistence in carbapenem-resistant Klebsiella pneumoniae. Nat Med. 05 2020;26(5):705–711. doi:10.1038/s41591-020-0825-4

41. Martens EC, Koropatkin NM, Smith TJ, Gordon JI. Complex glycan catabolism by the human gut microbiota: the Bacteroidetes Sus-like paradigm. J Biol Chem. Sep 2009;284(37):24673–7. doi:10.1074/jbc.R109.022848

42. Zhao S, Lieberman TD, Poyet M, et al. Adaptive Evolution within Gut Microbiomes of Healthy People. Cell Host Microbe. 05 2019;25(5):656–667.e8. doi:10.1016/j.chom.2019.03.007

43. Warsi OM, Andersson DI, Dykhuizen DE. Different adaptive strategies in E. coli populations evolving under macronutrient limitation and metal ion limitation. BMC Evol Biol. 05 18 2018;18(1):72. doi:10.1186/s12862-018-1191-4

44. Hickman RA, Munck C, Sommer MOA. Time-Resolved Tracking of Mutations Reveals Diverse Allele Dynamics during Escherichia coli Antimicrobial Adaptive Evolution to Single Drugs and Drug Pairs. Front Microbiol. 2017;8:893. doi:10.3389/fmicb.2017.00893

45. Auriol C, Bestel-Corre G, Claude JB, Soucaille P, Meynial-Salles I. Stress-induced evolution of Escherichia coli points to original concepts in respiratory cofactor selectivity. Proc Natl Acad Sci U S A. Jan 25 2011;108(4):1278–83. doi:10.1073/pnas.1010431108

46. Babel H, Krömer JO. Evolutionary engineering of E. coli MG1655 for tolerance against isoprenol. Biotechnol Biofuels. Nov 09 2020;13(1):183. doi:10.1186/s13068-020-01825-6

47. Juers DH, Matthews BW, Huber RE. LacZ β-galactosidase: structure and function of an enzyme of historical and molecular biological importance. Protein Sci. Dec 2012;21(12):1792–807. doi:10.1002/pro.2165

48. Rogers AWL, Tsolis RM, Bäumler AJ. Salmonella versus the Microbiome. Microbiol Mol Biol Rev. 02 17 2021;85(1)doi:10.1128/MMBR.00027-19

49. Hughes ER, Winter MG, Duerkop BA, et al. Microbial Respiration and Formate Oxidation as Metabolic Signatures of Inflammation-Associated Dysbiosis. Cell Host Microbe. Feb 08 2017;21(2):208–219. doi:10.1016/j.chom.2017.01.005

50. Gupta S, Allen-Vercoe E, Petrof EO. Fecal microbiota transplantation: in perspective. Therap Adv Gastroenterol. Mar 2016;9(2):229–39. doi:10.1177/1756283X15607414

51. Ianiro G, Murri R, Sciumè GD, et al. Incidence of Bloodstream Infections, Length of Hospital Stay, and Survival in Patients With Recurrent Clostridioides difficile Infection Treated With Fecal Microbiota Transplantation or Antibiotics: A Prospective Cohort Study. Ann Intern Med. 11 2019;171(10):695–702. doi:10.7326/M18-3635

52. Wortelboer K, Nieuwdorp M, Herrema H. Fecal microbiota transplantation beyond Clostridioides difficile infections. EBioMedicine. Jun 2019;44:716–729. doi:10.1016/j.ebiom.2019.05.066

53. Lee SM, Donaldson GP, Mikulski Z, Boyajian S, Ley K, Mazmanian SK. Bacterial colonization factors control specificity and stability of the gut microbiota. Nature. Sep 19 2013;501(7467):426–9. doi:10.1038/nature12447

54. Whitaker WR, Shepherd ES, Sonnenburg JL. Tunable Expression Tools Enable Single-Cell Strain Distinction in the Gut Microbiome. Cell. 04 20 2017;169(3):538–546.e12. doi:10.1016/j.cell.2017.03.041

55. Martinson JNV, Pinkham NV, Peters GW, et al. Rethinking gut microbiome residency and the Enterobacteriaceae in healthy human adults. ISME J. 09 2019;13(9):2306–2318. doi:10.1038/s41396-019-0435-7

56. Tyakht AV, Manolov AI, Kanygina AV, et al. Genetic diversity of Escherichia coli in gut microbiota of patients with Crohn’s disease discovered using metagenomic and genomic analyses. BMC Genomics. Dec 27 2018;19(1):968. doi:10.1186/s12864-018-5306-5

57. Woyke T, Doud DFR, Schulz F. The trajectory of microbial single-cell sequencing. Nat Methods. Oct 31 2017;14(11):1045–1054. doi:10.1038/nmeth.4469

58. Prosperi MC, Salemi M. QuRe: software for viral quasispecies reconstruction from next-generation sequencing data. Bioinformatics. Jan 2012;28(1):132–3. doi:10.1093/bioinformatics/btr627

59. Domingo E, Perales C. Viral quasispecies. PLoS Genet. 10 2019;15(10):e1008271. doi:10.1371/journal.pgen.1008271

60. Yamada C, Gotoh A, Sakanaka M, et al. Molecular Insight into Evolution of Symbiosis between Breast-Fed Infants and a Member of the Human Gut Microbiome Bifidobacterium longum. Cell Chem Biol. Apr 2017;24(4):515–524.e5. doi:10.1016/j.chembiol.2017.03.012

61. Zerbino DR, Birney E. Velvet: algorithms for de novo short read assembly using de Bruijn graphs. Genome Res. May 2008;18(5):821–9. doi:10.1101/gr.074492.107

62. Gao S, Bertrand D, Chia BK, Nagarajan N. OPERA-LG: efficient and exact scaffolding of large, repeat-rich eukaryotic genomes with performance guarantees. Genome Biol. May 2016;17:102. doi:10.1186/s13059-016-0951-y

63. Gao S, Bertrand D, Nagarajan N. FinIS: Improved in silico Finishing Using an Exact Quadratic Programming Formulation. Lect Notes Comput Sci. 2012;7534:314–325.

64. Li H. Aligning sequence reads, clone sequences and assembly contigs with BWA-MEM. arXiv. 2013;(1303.3997v2)

65. Segata N, Waldron L, Ballarini A, Narasimhan V, Jousson O, Huttenhower C. Metagenomic microbial community profiling using unique clade-specific marker genes. Nat Methods. Jun 2012;9(8):811–4. doi:10.1038/nmeth.2066

66. Franzosa EA, McIver LJ, Rahnavard G, et al. Species-level functional profiling of metagenomes and metatranscriptomes. Nat Methods. 11 2018;15(11):962–968. doi:10.1038/s41592-018-0176-y

67. Salter SJ, Cox MJ, Turek EM, et al. Reagent and laboratory contamination can critically impact sequence-based microbiome analyses. BMC Biol. Nov 12 2014;12:87. doi:10.1186/s12915-014-0087-z

68. Segata N, Izard J, Waldron L, et al. Metagenomic biomarker discovery and explanation. Genome Biol. Jun 2011;12(6):R60. doi:10.1186/gb-2011-12-6-r60

69. Hawinkel S, Mattiello F, Bijnens L, Thas O. A broken promise: microbiome differential abundance methods do not control the false discovery rate. Brief Bioinform. 01 18 2019;20(1):210–221. doi:10.1093/bib/bbx104

70. Morton JT, Marotz C, Washburne A, et al. Establishing microbial composition measurement standards with reference frames. Nat Commun. 06 20 2019;10(1):2719. doi:10.1038/s41467-019-10656-5

71. Inouye M, Dashnow H, Raven LA, et al. SRST2: Rapid genomic surveillance for public health and hospital microbiology labs. Genome Med. 2014;6(11):90. doi:10.1186/s13073-014-0090-6

72. Alcock BP, Raphenya AR, Lau TTY, et al. CARD 2020: antibiotic resistome surveillance with the comprehensive antibiotic resistance database. Nucleic Acids Res. 01 08 2020;48(D1):D517–D525. doi:10.1093/nar/gkz935

73. Kurtz S, Phillippy A, Delcher AL, et al. Versatile and open software for comparing large genomes. Genome Biol. 2004;5(2):R12. doi:10.1186/gb-2004-5-2-r12

74. Wilm A, Aw PP, Bertrand D, et al. LoFreq: a sequence-quality aware, ultra-sensitive variant caller for uncovering cell-population heterogeneity from high-throughput sequencing datasets. Nucleic Acids Res. Dec 2012;40(22):11189–201. doi:10.1093/nar/gks918

75. Hinrichs AS, Karolchik D, Baertsch R, et al. The UCSC Genome Browser Database: update 2006. Nucleic Acids Res. Jan 01 2006;34(Database issue):D590–8. doi:10.1093/nar/gkj144

76. Pracana R, Priyam A, Levantis I, Nichols RA, Wurm Y. The fire ant social chromosome supergene variant Sb shows low diversity but high divergence from SB. Mol Ecol. Jun 2017;26(11):2864–2879. doi:10.1111/mec.14054

77. Quinlan AR. BEDTools: The Swiss-Army Tool for Genome Feature Analysis. Curr Protoc Bioinformatics. Sep 2014;47:11.12.1-34. doi:10.1002/0471250953.bi1112s47

78. Spedicato G. Discrete Time Markov Chains with R. The R Journal. 2017;

79. Cingolani P, Platts A, Wang lL, et al. A program for annotating and predicting the effects of single nucleotide polymorphisms, SnpEff: SNPs in the genome of Drosophila melanogaster strain w1118; iso-2; iso-3. Fly (Austin). 2012 Apr-Jun 2012;6(2):80–92. doi:10.4161/fly.19695

80. Hahsler M, Piekenbrock M, Doran D. dbscan: Fast Density-Based Clustering with R. Journal of Statistical Software. 2019;91:1–30.

81. Galata V, Fehlmann T, Backes C, Keller A. PLSDB: a resource of complete bacterial plasmids. Nucleic Acids Res. 01 2019;47(D1):D195–D202. doi:10.1093/nar/gky1050

82. Ondov BD, Treangen TJ, Melsted P, et al. Mash: fast genome and metagenome distance estimation using MinHash. Genome Biol. 06 2016;17(1):132. doi:10.1186/s13059-016-0997-x

83. Quan S, Ray JC, Kwota Z, et al. Adaptive evolution of the lactose utilization network in experimentally evolved populations of Escherichia coli. PLoS Genet. Jan 2012;8(1):e1002444. doi:10.1371/journal.pgen.1002444

84. Tsuchido T, VanBogelen RA, Neidhardt FC. Heat shock response in Escherichia coli influences cell division. Proc Natl Acad Sci U S A. Sep 1986;83(18):6959–63. doi:10.1073/pnas.83.18.6959

85. Trubetskoy D, Proux F, Allemand F, Dreyfus M, Iost I. SrmB, a DEAD-box helicase involved in Escherichia coli ribosome assembly, is specifically targeted to 23S rRNA in vivo. Nucleic Acids Res. Oct 2009;37(19):6540–9. doi:10.1093/nar/gkp685

86. Garoff L, Huseby DL, Praski Alzrigat L, Hughes D. Effect of aminoacyl-tRNA synthetase mutations on susceptibility to ciprofloxacin in Escherichia coli. J Antimicrob Chemother. 12 01 2018;73(12):3285–3292. doi:10.1093/jac/dky356

87. Aponte RA, Zimmermann S, Reinstein J. Directed evolution of the DnaK chaperone: mutations in the lid domain result in enhanced chaperone activity. J Mol Biol. May 28 2010;399(1):154–67. doi:10.1016/j.jmb.2010.03.060

88. Mundhada H, Seoane JM, Schneider K, et al. Increased production of L-serine in Escherichia coli through Adaptive Laboratory Evolution. Metab Eng. 01 2017;39:141–150. doi:10.1016/j.ymben.2016.11.008

89. Conrad TM, Frazier M, Joyce AR, et al. RNA polymerase mutants found through adaptive evolution reprogram Escherichia coli for optimal growth in minimal media. Proc Natl Acad Sci U S A. Nov 23 2010;107(47):20500–5. doi:10.1073/pnas.0911253107

90. Li Y, Powell DA, Shaffer SA, et al. LPS remodeling is an evolved survival strategy for bacteria. Proc Natl Acad Sci U S A. May 29 2012;109(22):8716–21. doi:10.1073/pnas.1202908109

