## Supplementary figures and tables for "Long-term ecological and evolutionary dynamics in the gut microbiomes of carbapenemase-producing *Enterobacteriaceae* colonized subjects"

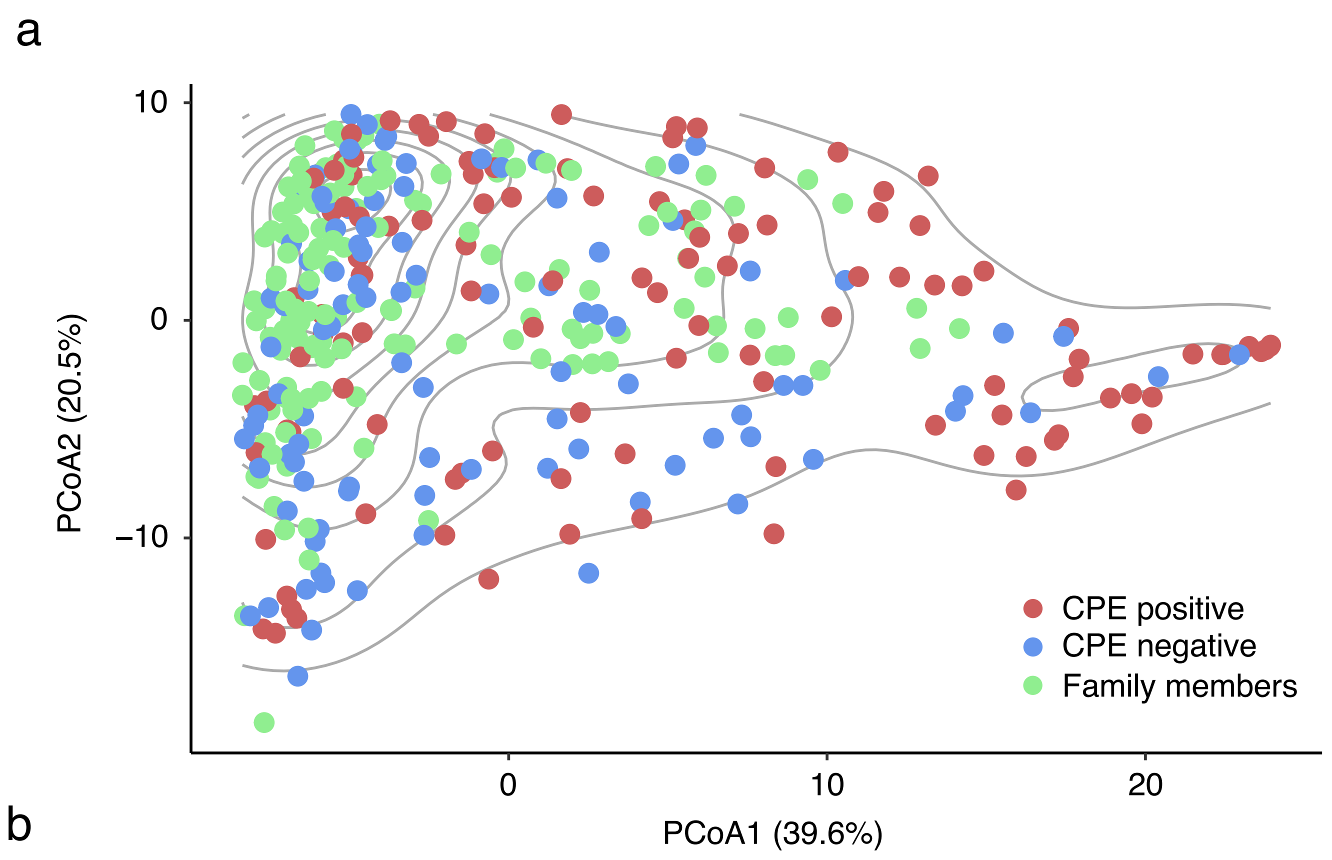


**<u>Supplementary Figure 1:</u>** (a) Principal coordinates analysis (PCoA) plot similar to **Figure 1a** but calculated based on genus-level weighted UniFrac distance. Note that while the location of points is flipped with respect to the x-axis for **Figure 1a**, the general distribution of points is similar with regions of high density on the left having more family members while a less dense cluster on the right has more CPE positive points. (b) Dendrogram for hierarchical clustering based on distance in PCoA1-PCoA2 space in **Figure 1a**, and number of clusters determined by AIC. The labels I–IV correspond to those in **Figure 1a**. Dendrogram tips are labeled with the library ID of each sample (see **Supplementary File 1**).

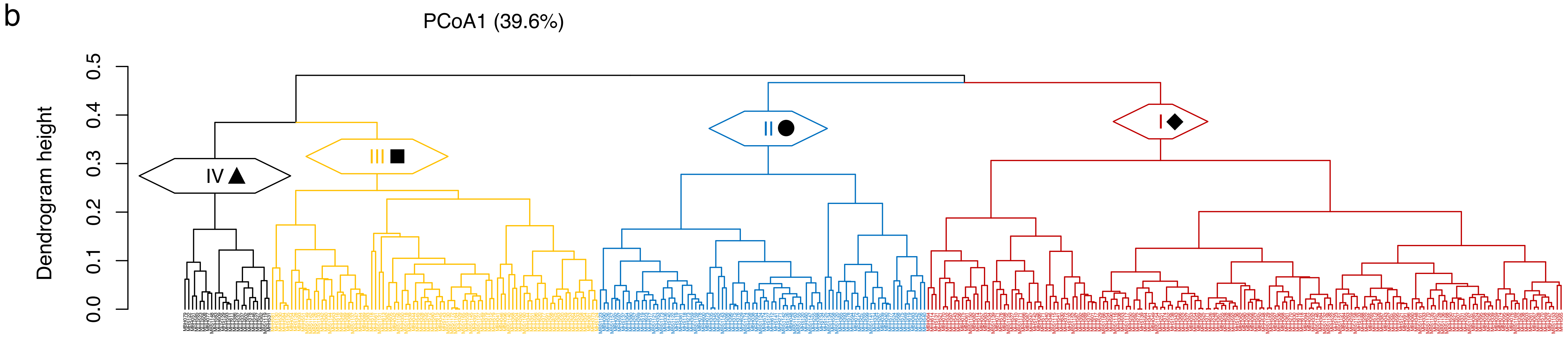

a

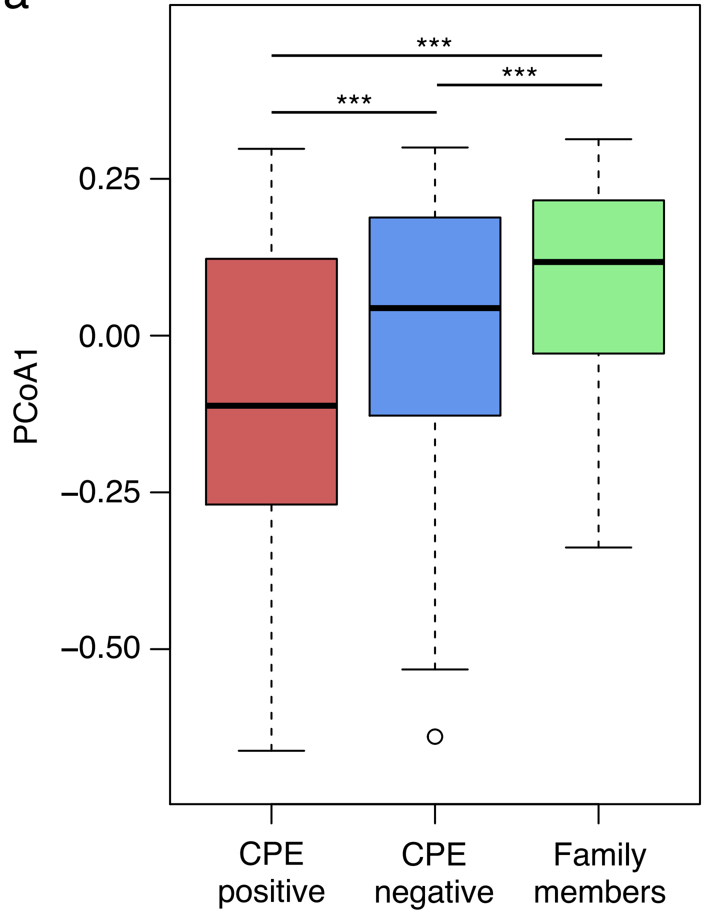


b

| Genus | Sign | Correlation |
|---|---|---|
| *Bacteroides* | + | 0.736 |
| *Escherichia* | – | 0.722 |
| *Pantoea* | – | 0.543 |
| *Klebsiella* | – | 0.354 |
| Lachnospiraceae unclassified | + | 0.285 |
| *Blautia* | + | 0.260 |
| *Oscillibacter* | + | 0.249 |
| *Scardovia* | – | 0.243 |
| *Sutterella* | + | 0.241 |
| *Odoribacter* | + | 0.240 |
| *Holdemania* | + | 0.230 |
| *Eubacterium* | + | 0.216 |

**Supplementary Figure 2:** (a) Comparison of principal coordinates analysis (PCoA) values (**Figure 1a**) across different groups of samples based on the first principal coordinate (PCoA1). Wilcoxon rank-sum $p<0.01$ for all pairwise comparisons. (b) Genera most correlated with PCoA1 (**Figure 1a**). Sign (–): Negative correlation with PCoA1 (most abundant in configuration IV samples). Sign (+): Positive correlation with PCoA1 (least abundant in configuration IV samples).

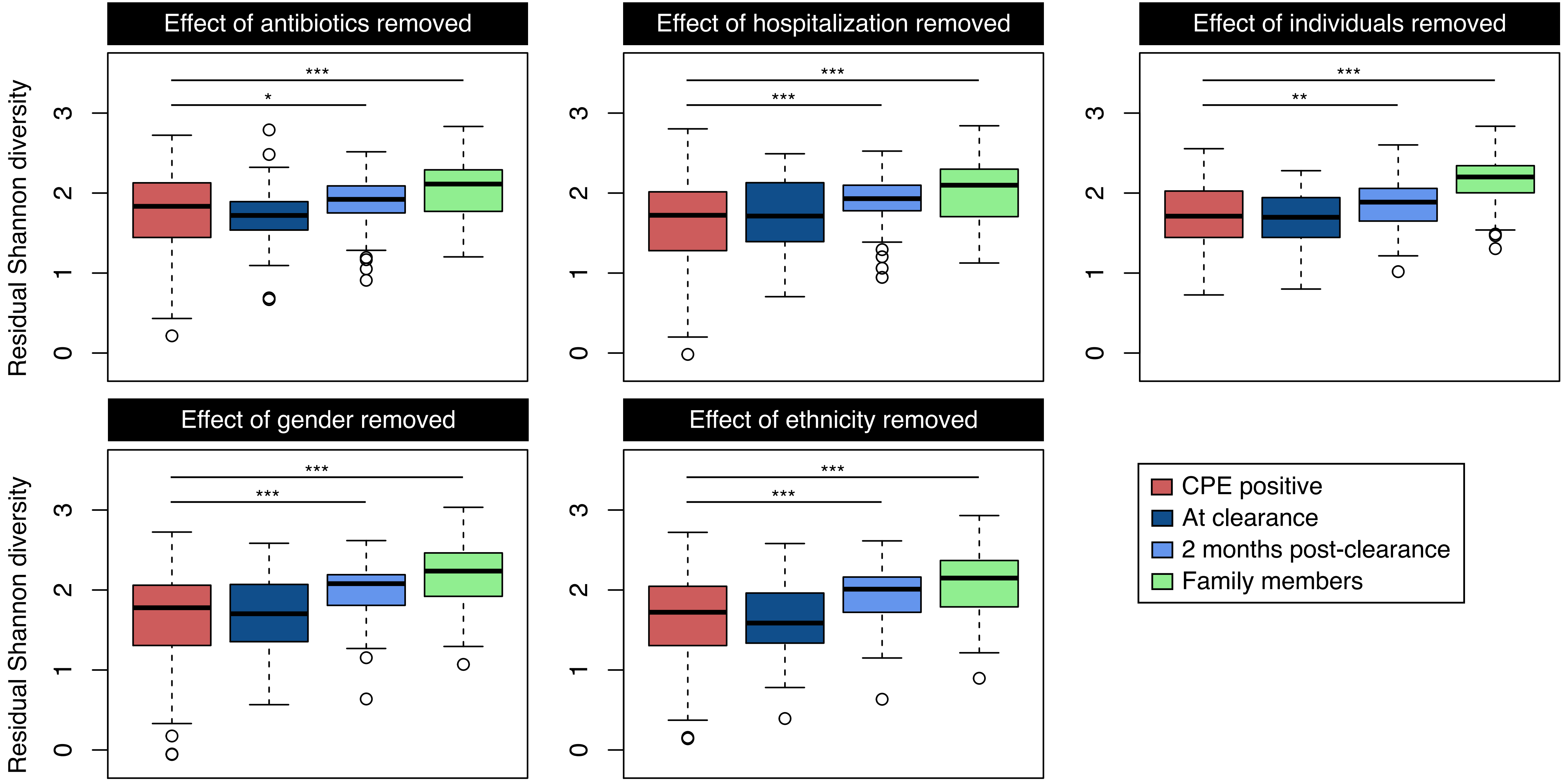


**Supplementary Figure 3:** Residual Shannon diversity, after subtracting the intercept term due to the random effect in a linear mixed-effects model that treats colonization status as the fixed effect, and the stated covariates as random effects (i.e. antibiotic usage since last visit, hospitalization status, individual subjects, gender and ethnicity; see **Supplementary File 1**). The p-values originate as part of the output from the respective linear mixed effect models, comparing CPE positive timepoints against other groups. *** = $p<0.01$, ** = $p<0.05$, * = $p<0.1$.

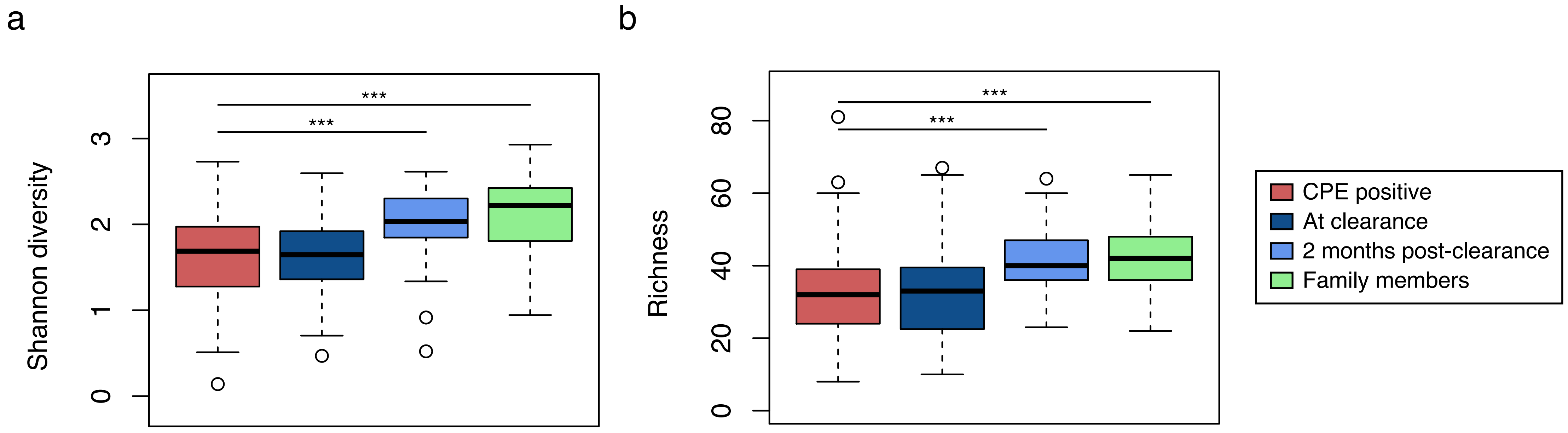


**Supplementary Figure 4:** (a) Boxplots showing genus-level Shannon diversity distributions for different timepoints for index patients ("CPE positive" = during colonization, "At clearance" = within 1 month of decolonization, and "2 months post-clearance" = time points that were >2 months after decolonization), and all timepoints for family members. This plot is similar to **Figure 1a** but with all *Enterobacteriaceae* counts removed and relative abundances renormalized to 1. (b) Corresponding microbial richness values. *** = Wilcoxon rank-sum p<0.01 and all other comparisons were not statistically significant.

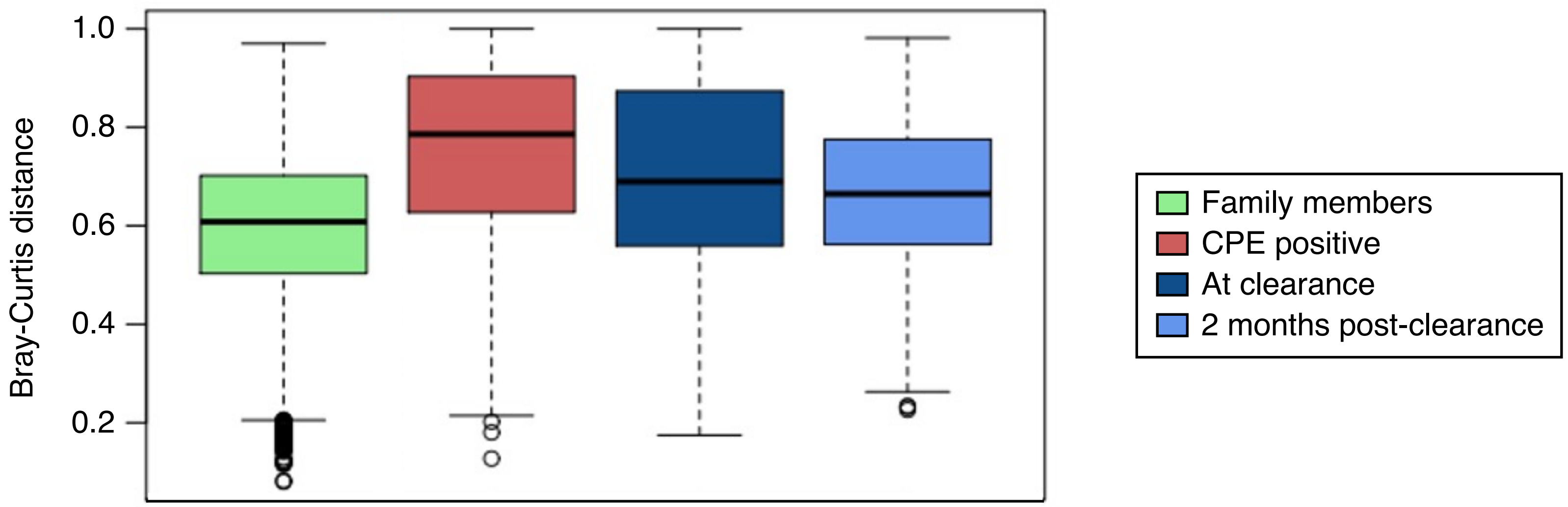


**<u>Supplementary Figure 5:</u>** Boxplots quantifying the distributions of pairwise genus-level Bray-Curtis distances. Each pair consists of one sample from a family member, and a second sample from one of the following groups: family members, CPE positive and CPE negative at point of clearance, as well as more than two months post-clearance. Wilcoxon rank-sum $p<0.01$ for all pairwise comparisons with the family member group.

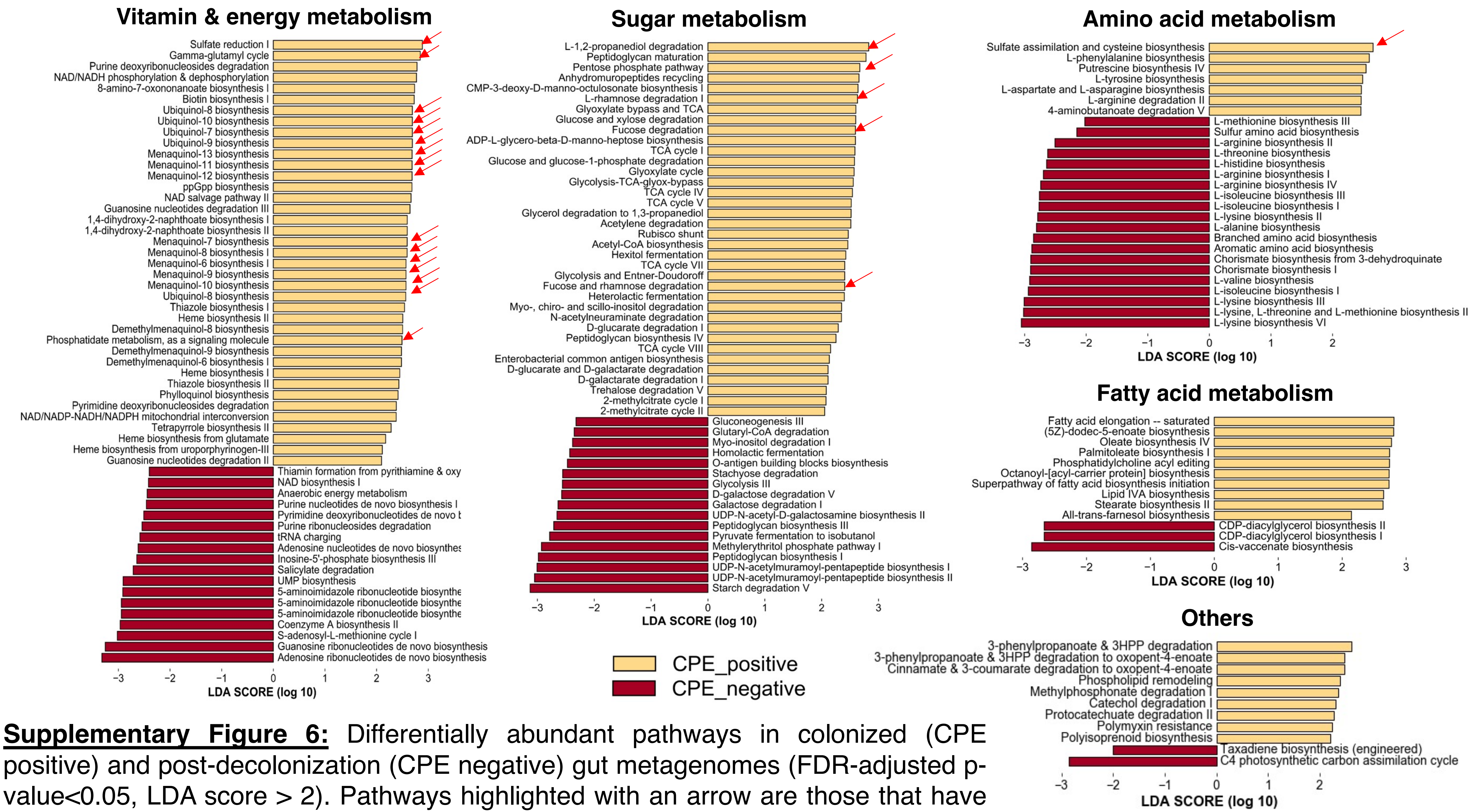


**Supplementary Figure 6:** Differentially abundant pathways in colonized (CPE positive) and post-decolonization (CPE negative) gut metagenomes (FDR-adjusted p-value<0.05, LDA score > 2). Pathways highlighted with an arrow are those that have potential associations with gut inflammation and aerobic respiration.

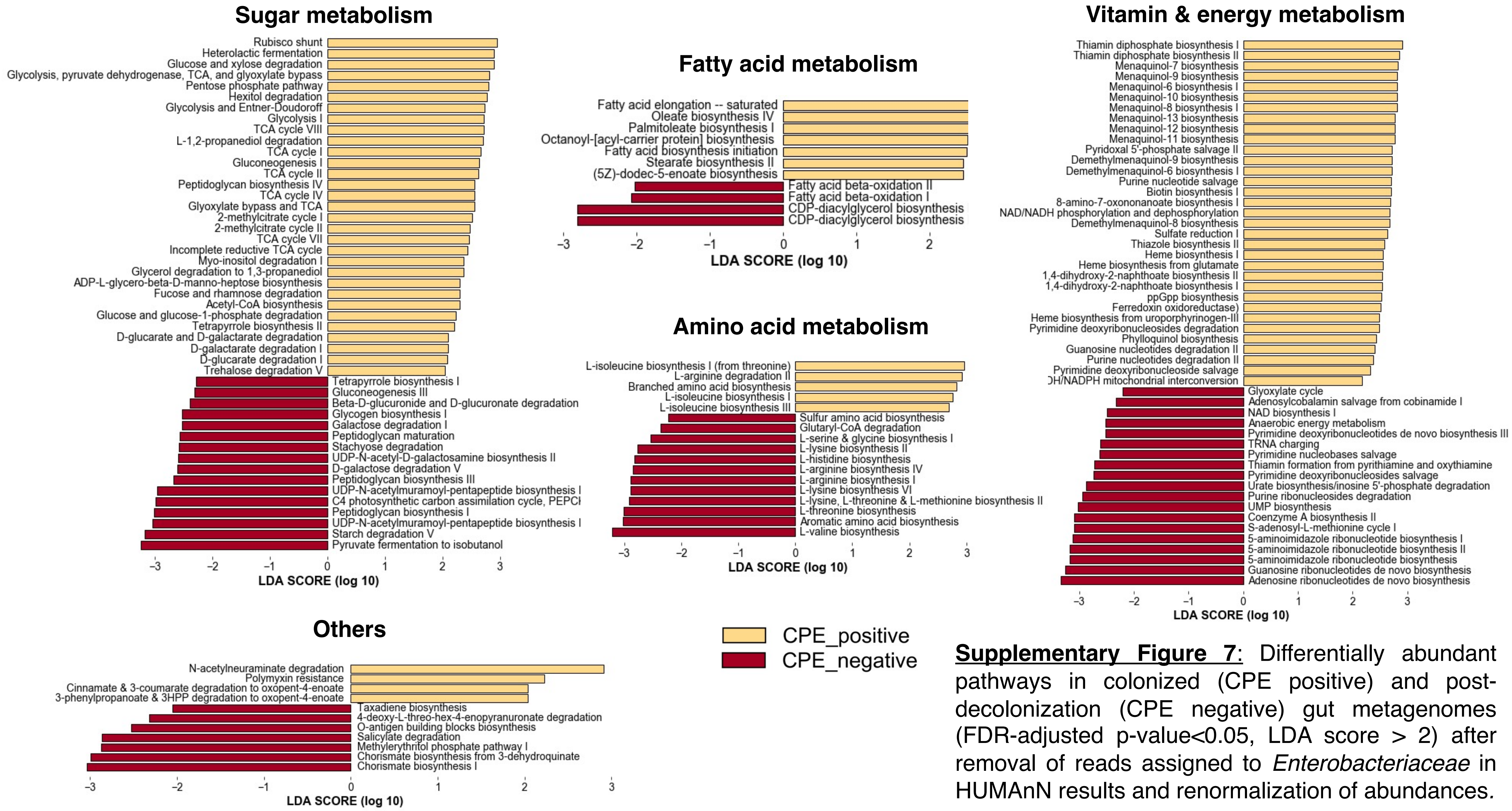


**<u>Supplementary Figure 7</u>**: Differentially abundant pathways in colonized (CPE positive) and post-decolonization (CPE negative) gut metagenomes (FDR-adjusted p-value<0.05, LDA score > 2) after removal of reads assigned to *Enterobacteriaceae* in HUMAnN results and renormalization of abundances.

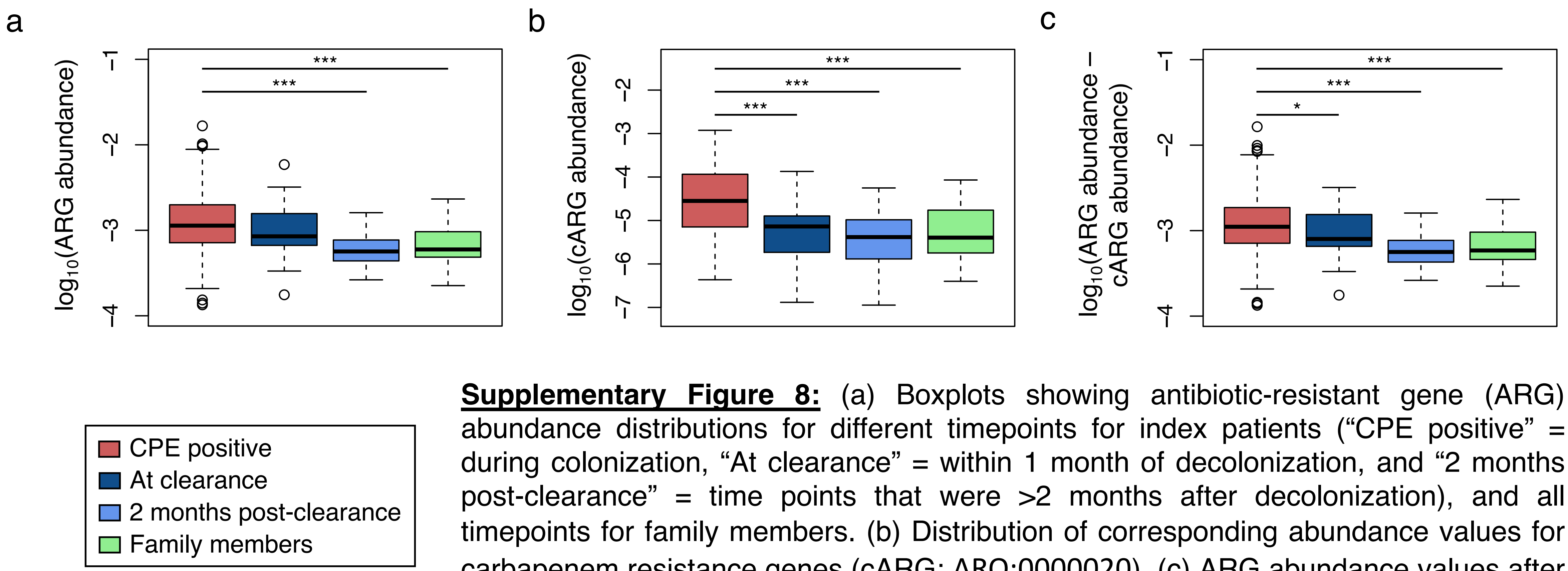


**Supplementary Figure 8:** (a) Boxplots showing antibiotic-resistant gene (ARG) abundance distributions for different timepoints for index patients (“CPE positive” = during colonization, “At clearance” = within 1 month of decolonization, and “2 months post-clearance” = time points that were >2 months after decolonization), and all timepoints for family members. (b) Distribution of corresponding abundance values for carbapenem resistance genes (cARG; ARO:0000020). (c) ARG abundance values after subtracting cARG abundances. *** = Wilcoxon rank-sum $p<0.01$, * = Wilcoxon rank-sum $p<0.1$, and all other comparisons were not statistically significant.

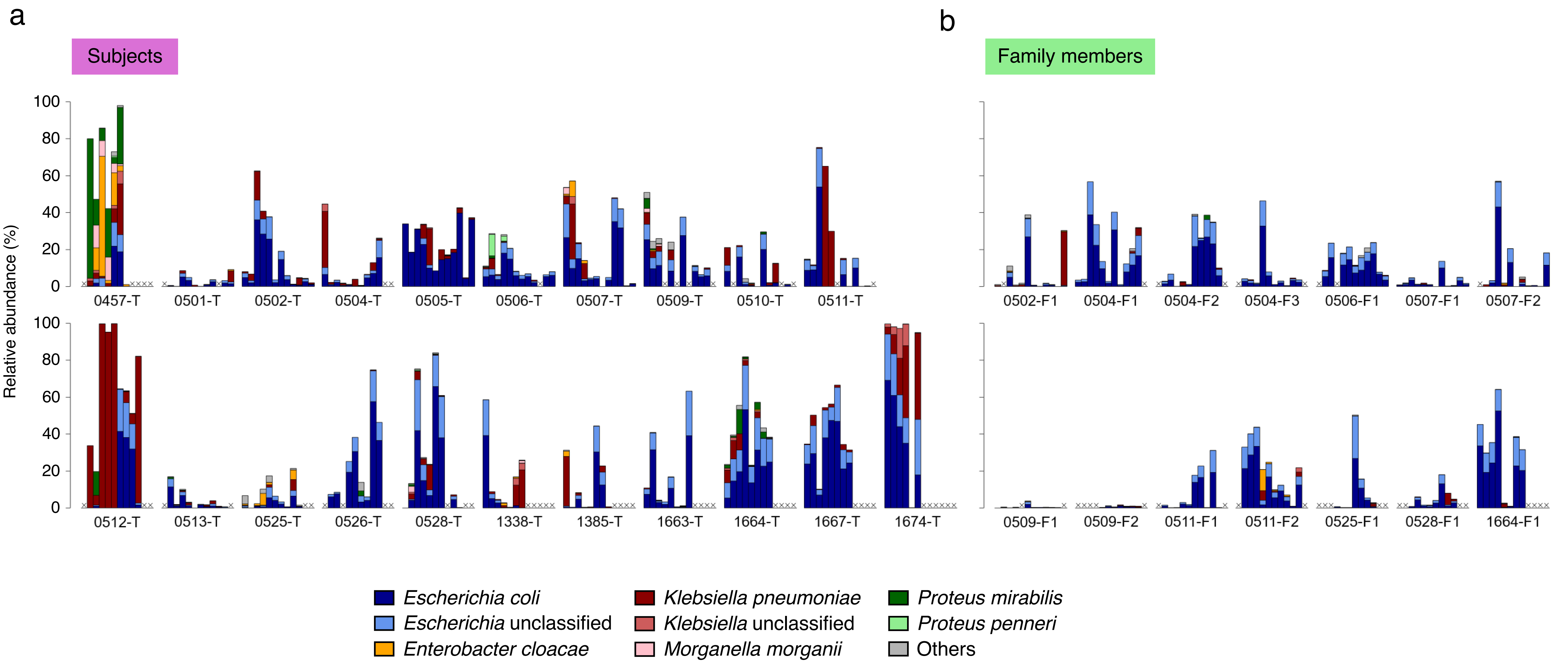


**Supplementary Figure 9:** Bar plots showing the relative abundances of various Enterobacteriaceae species in (a) subjects and (b) family members, over the course of sample collection. Each sub-panel consists of 12 bars, representing consecutively visit 0 (V00) to visit 11 (V11). An "X" at a particular position indicates that a sample corresponding to that visit number was not collected. Only individuals with samples collected at six or more visits are shown.

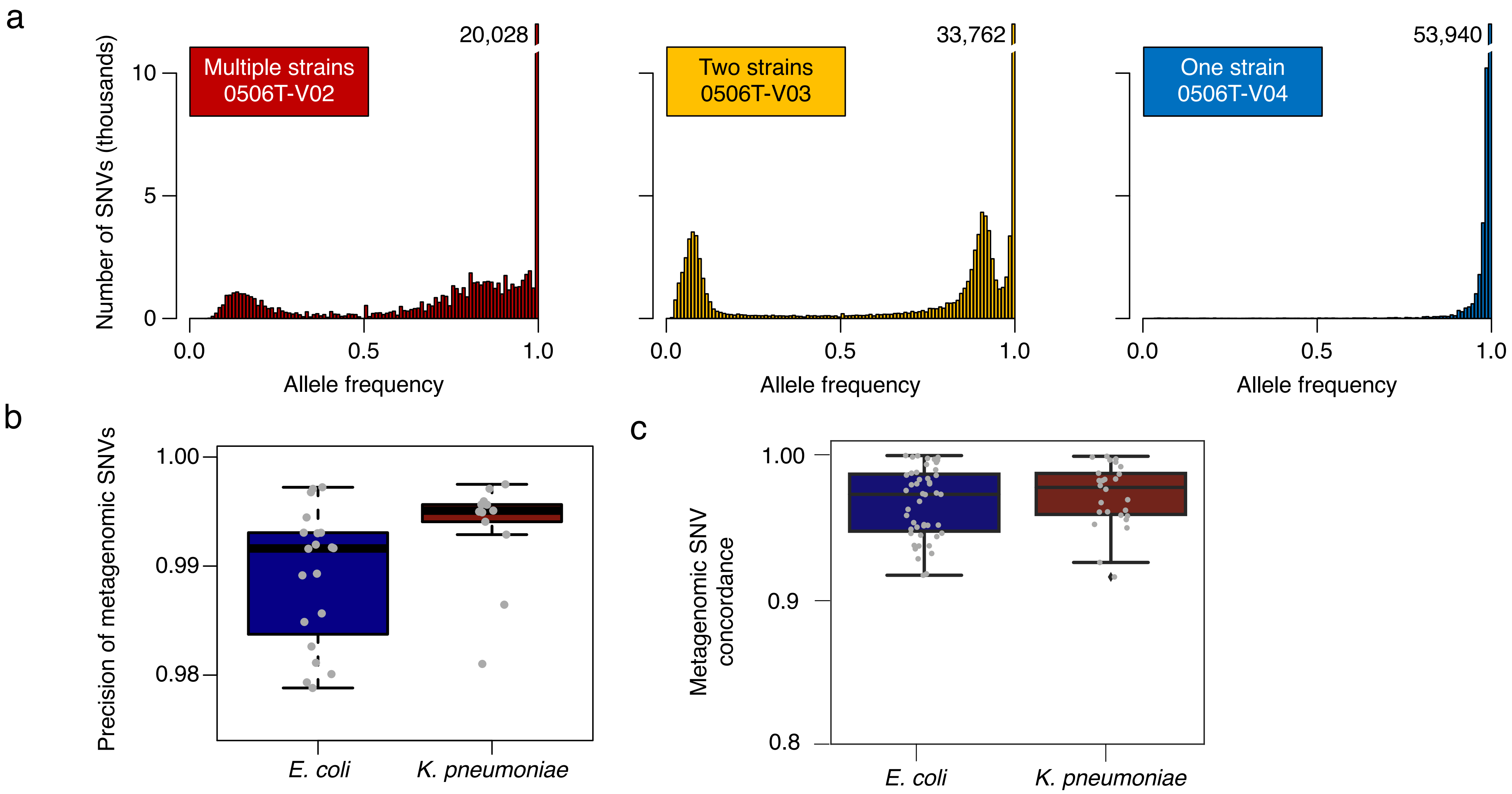


**Supplementary Figure 10:** (a) Representative allele frequency spectra for samples classified as "one strain", "two strains" and "multiple strains", obtained from 3 consecutive timepoints in subject 0506-T. (b) Boxplots depicting the precision of metagenomic SNVs (One strain, allele frequency ≥0.98) evaluated using SNVs present in corresponding CPE isolates where available. (c) Fraction of metagenomic SNVs (all allele frequencies) called using a shared species reference that were recapitulated using sample-specific analysis with CPE isolate references.

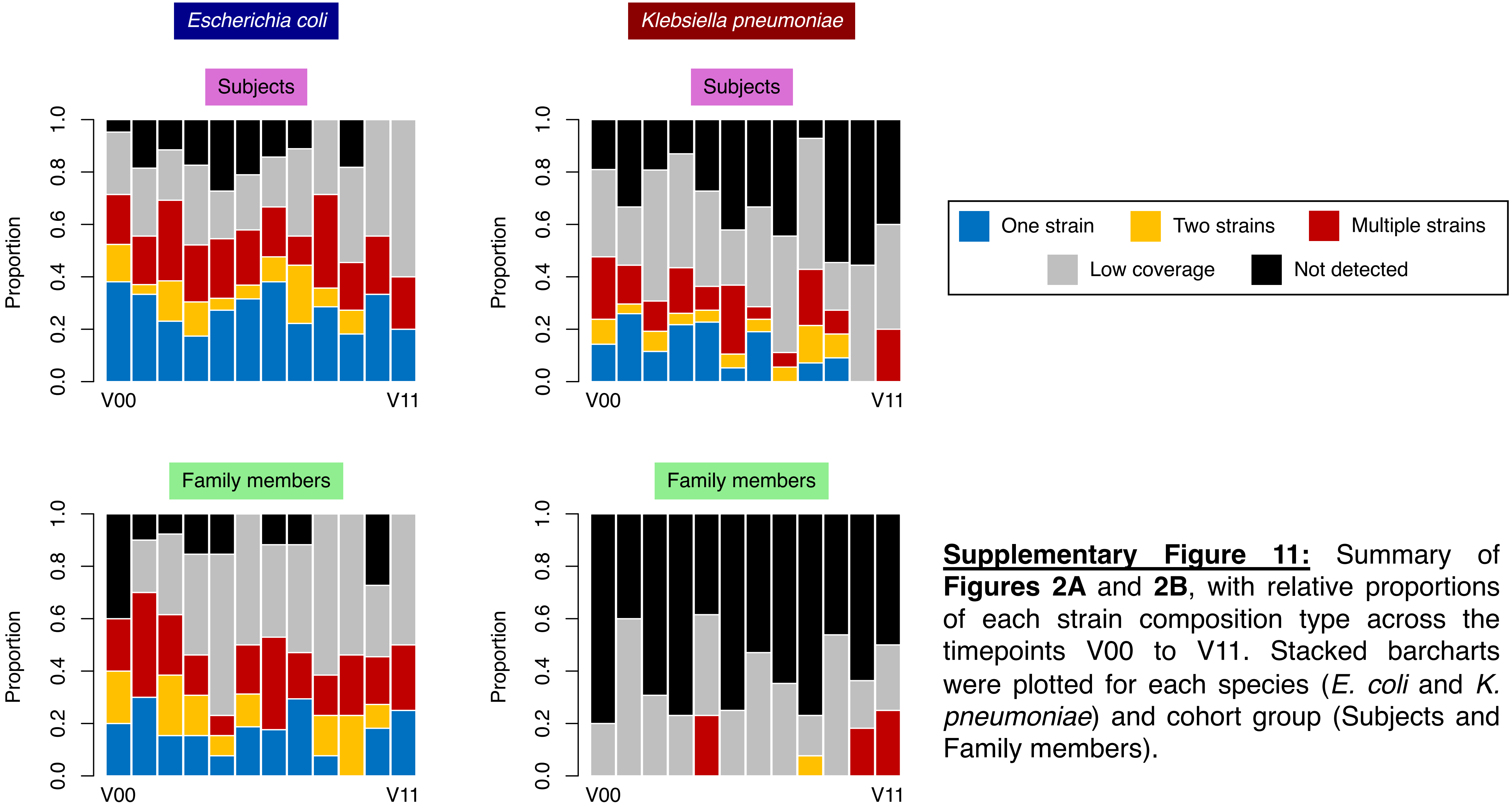


**Supplementary Figure 11:** Summary of **Figures 2A** and **2B**, with relative proportions of each strain composition type across the timepoints V00 to V11. Stacked barcharts were plotted for each species (*E. coli* and *K. pneumoniae*) and cohort group (Subjects and Family members).

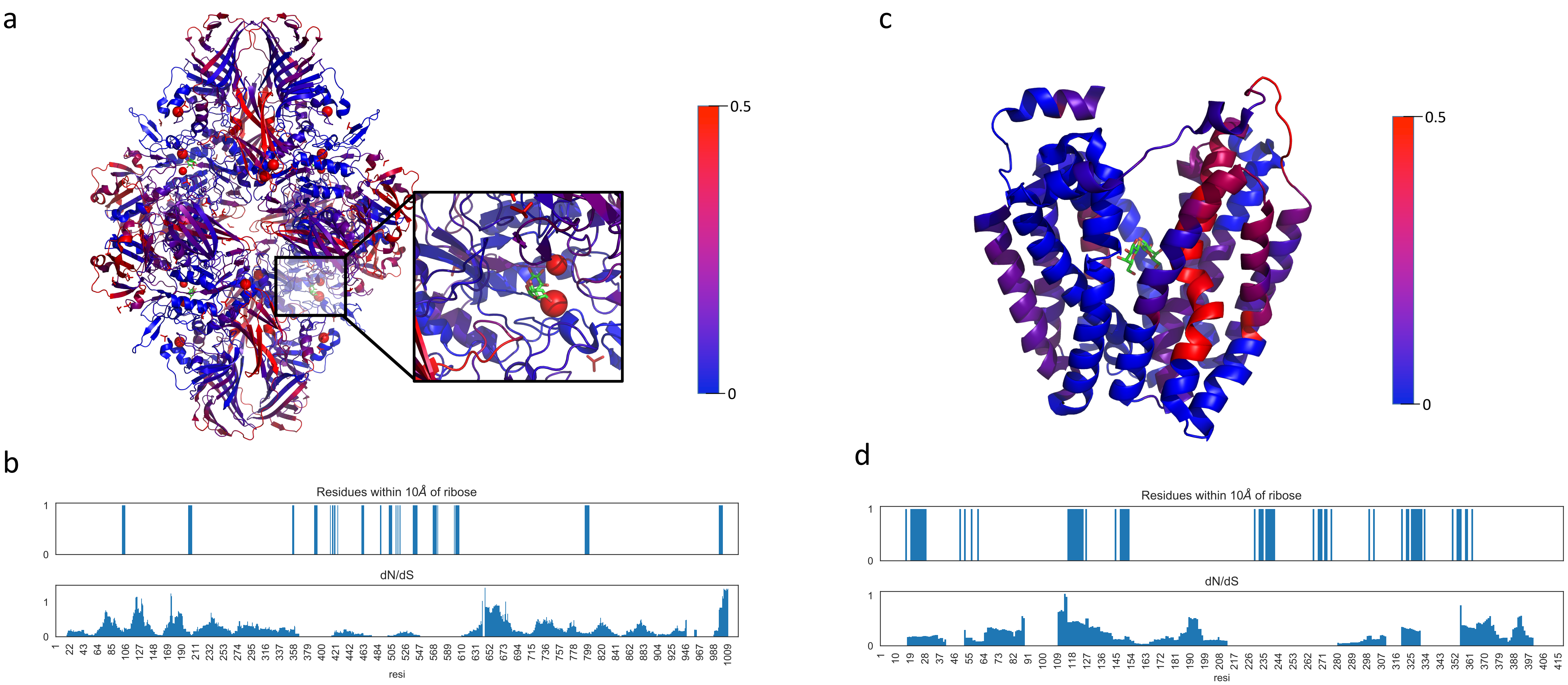


**Supplementary Figure 12**: (a) Visualization of dN/dS rates for the lacZ tetramer (4dux) using a 30aa window based on metagenomics-derived SNVs (**Table 2**). The inset shows a close-up view of lacZ's sugar binding active site. In general, dN/dS rates were lower for lacZ in regions close to the active site (Wilcoxon p-value<$2.6\times10^{-9}$) except for around position 1000. (b) Plot of dN/dS values and regions of lacZ that are close to ribose. (c) Visualization of dN/dS rates for lacY (1pv7) using a 30aa window. Interestingly, lacY exhibited a distinct trend with a few regions of higher dN/dS rates close to the active site (Wilcoxon p-value<0.029). (d) Plot of dN/dS values and regions close to the lacY lactose symporter channel.

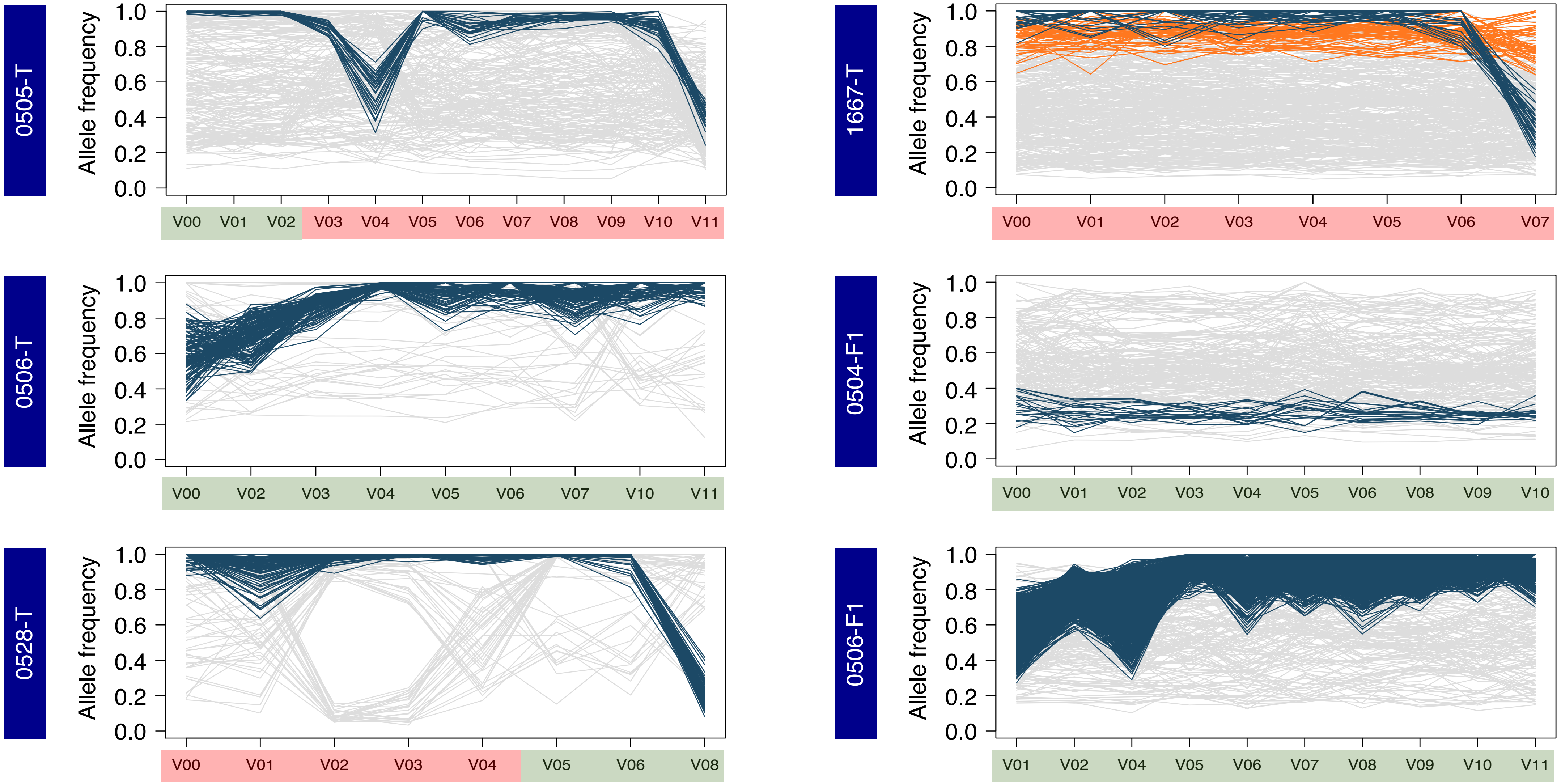


**Supplementary Figure 13:** Sub-strain variations over time for other individuals (*E. coli*). Selected figures were plotted for subjects with >3 consecutive single-strain timepoints. Colored lines depict clusters of SNVs that were identified. Timepoints are shaded based on whether they were determined to be CPE colonized (red) or not (green). Note that while a cluster of SNVs typically tends to be dominant in a subject across timepoints (colored in dark blue, except for 0504-F1), they can drop in frequency at some timepoints, and this does not necessarily coincide with the CPE decolonization event as was observed for 1674-T (e.g. V08 for 0528-T).

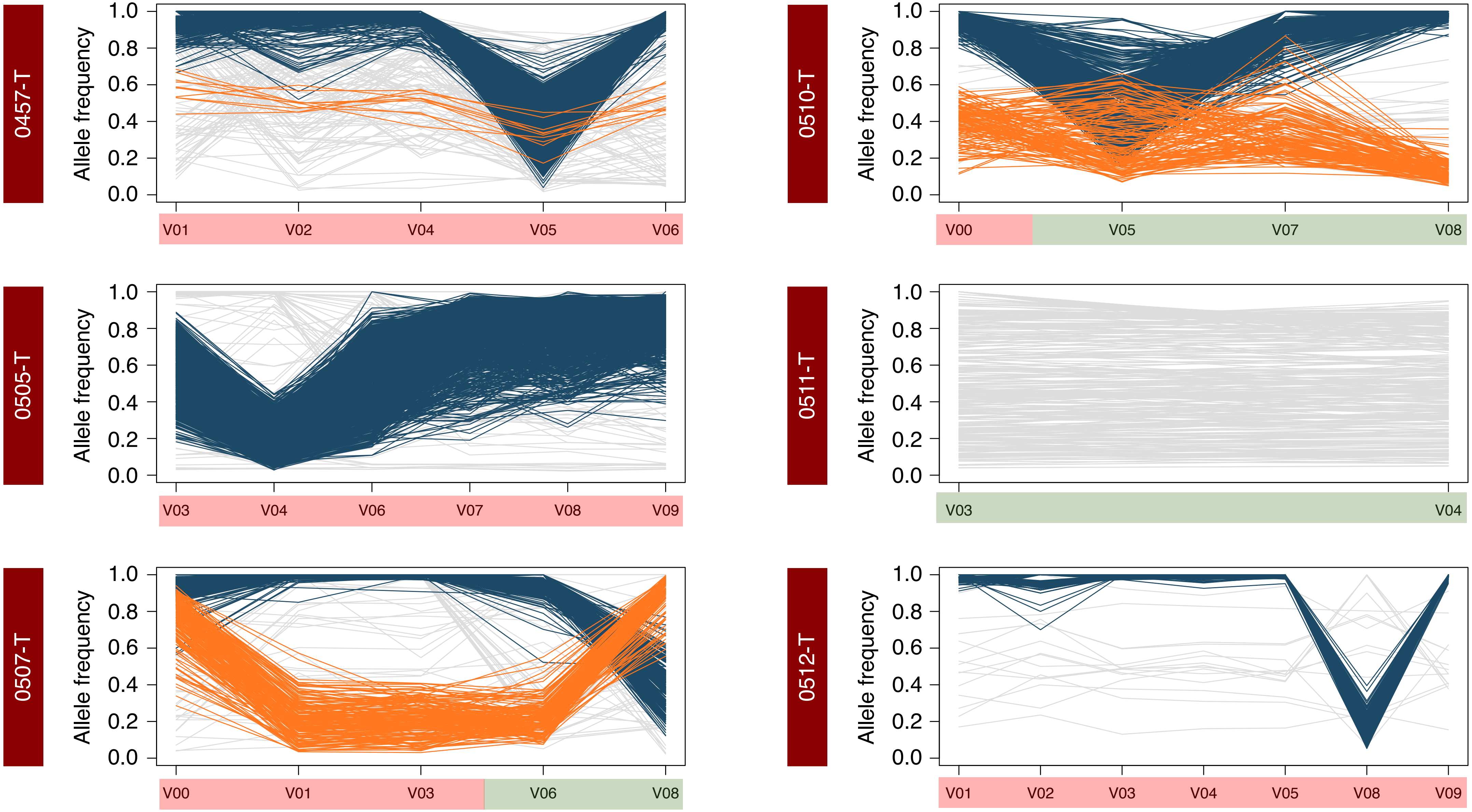


**<u>Supplementary Figure 14:</u>** Sub-strain variations over time for other individuals (*K. pneumoniae*). Figures were plotted for subjects with ≥2 single-strain timepoints. Colored lines depict clusters of SNVs that were identified. Timepoints are shaded based on whether they were determined to be CPE colonized (red) or not (green). Note that similar to the observation in **Supplementary Figure 13**, the dominant sub-strain in a subject can switch at timepoints that do not coincide with change in CPE colonization status (e.g. V01 and V08 for 0507-T).

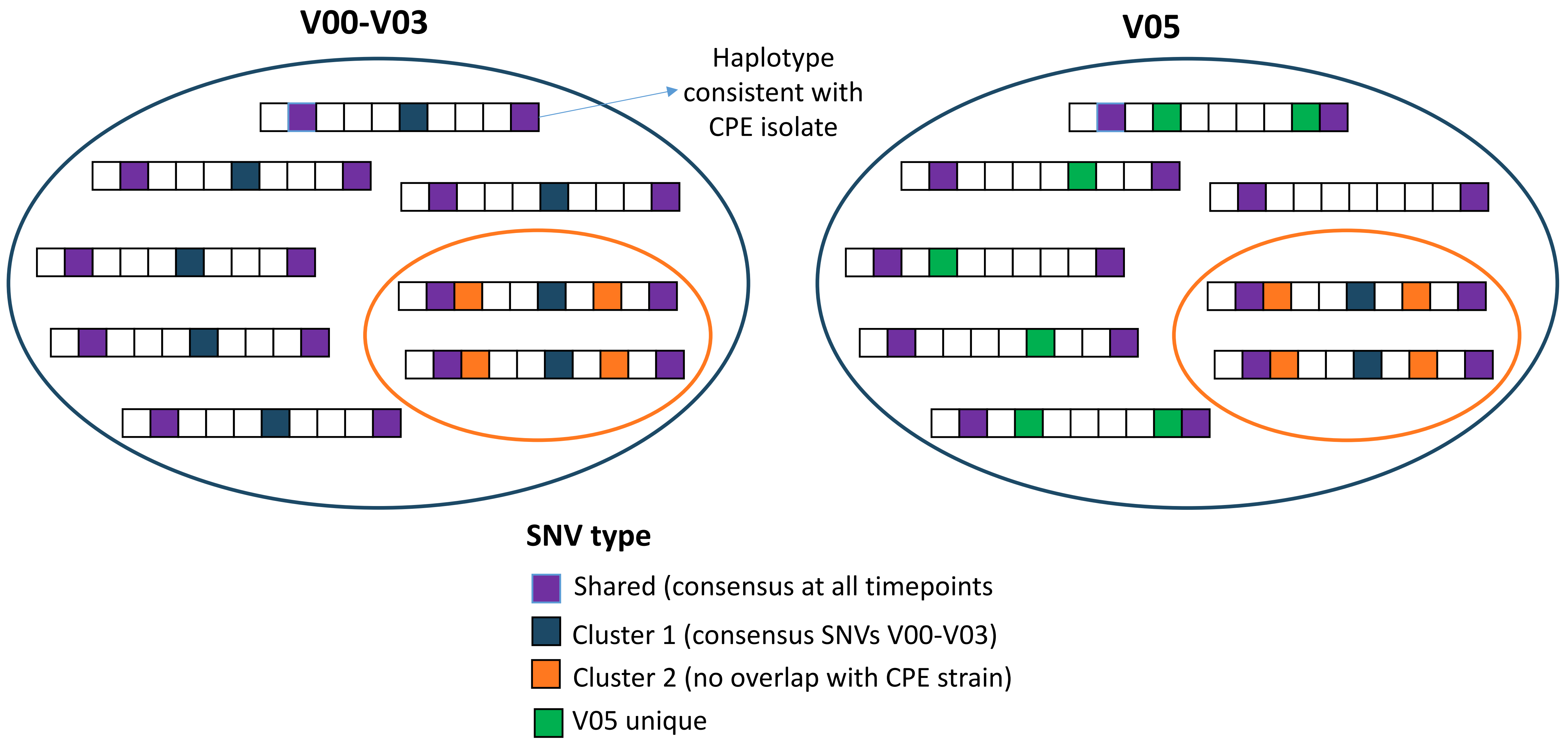


**Supplementary Figure 15:** Pictorial depiction of haplotype populations consistent with the sub-strain clusters seen in **Figure 3** for *E. coli*. Each strip of boxes represents one haplotype, with white boxes representing reference bases and colored bases representing SNVs that are part of different time-series clusters from **Figure 3a**. Note that the population on the left depicts the putative state for timepoints V00-V03 and at timepoint V05 there is a loss of the CPE sub-strain to give the population seen on the right.

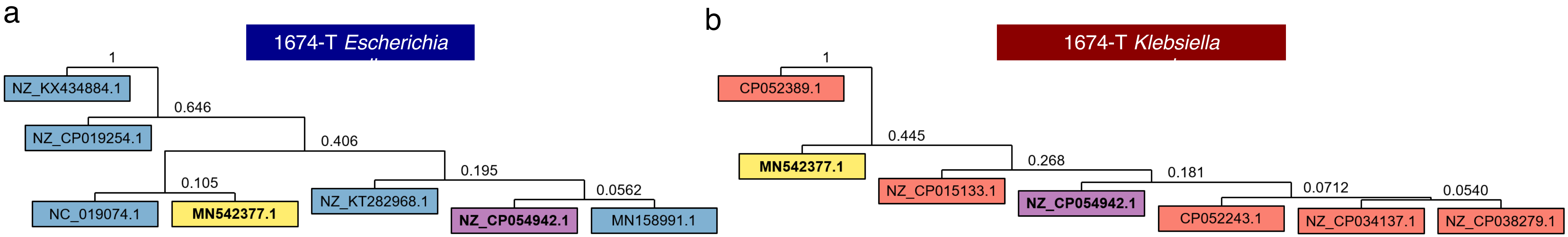


c

| Plasmid accession | Plasmid name | Plasmid length (bp) | *E. coli* | | | | *K. pneumoniae* | | | |
|---|---|---|---|---|---|---|---|---|---|---|
| | | | V00 | V01 | V02 | V03 | V00 | V01 | V02 | V03 |
| MN542377.1 | *K. pneumoniae* strain 494 plasmid pKPC2_sg1 | 71861 | 0.979 | 0.983 | 0.984 | 0.979 | 0.972 | 0.972 | 0.972 | 0.981 |
| NZ_CP054942.1 | *E. coli* strain MS6192 plasmid pMS6192B | 76661 | 0.828 | 0.824 | 0.809 | 0.853 | 0.862 | *0.049* | *0.049* | *0.068* |
| NZ_KX434884.1 | *K. pneumoniae* strain KP2442 plasmid pKP2442_4c285 | 12465 | 0.866 | 0.866 | 0.866 | 0.866 | *0.109* | *0.109* | *0.109* | *0.109* |
| NZ_CP019254.1 | *E. coli* strain 13KWH46 plasmid p13KWH46-4 | 76784 | 0.912 | 0.912 | 0.911 | 0.912 | - | - | - | - |
| NC_019074.1 | *E. coli* plasmid pHNDD81-1 | 11744 | 0.874 | 0.874 | 0.874 | 0.874 | *0.640* | *0.553* | *0.553* | *0.611* |
| NZ_KT282968.1 | *E. coli* strain EC012 plasmid pEC012 | 139622 | 0.891 | 0.888 | 0.884 | 0.884 | *0.074* | *0.069* | *0.067* | *0.102* |
| MN158991.1 | *E. coli* strain TREC8 plasmid pTREC8 | 118200 | 0.852 | 0.860 | 0.842 | 0.869 | *0.412* | *0.103* | *0.103* | *0.120* |
| CP052389.1 | *K. pneumoniae* strain C17KP0052 plasmid pC17KP0052-1 | 242988 | - | - | - | - | 0.997 | 0.997 | 0.997 | 0.997 |
| NZ_CP015133.1 | *K. pneumoniae* strain Kpn555 plasmid pKPN-d6b | 26450 | - | - | - | - | 0.907 | 0.907 | 0.907 | 0.993 |
| CP052243.1 | *K. pneumoniae* strain E17KP0019 plasmid pE17KP0019-2 | 246731 | *0.106* | *0.107* | *0.108* | *0.107* | 0.923 | 0.898 | 0.900 | 0.923 |
| NZ_CP034137.1 | *K. quasipneumoniae* strain G747 plasmid pG747_150.8Kb | 150833 | *0.176* | *0.180* | *0.180* | *0.180* | 0.869 | 0.853 | 0.858 | 0.896 |
| NZ_CP038279.1 | *R. ornithinolytica* strain WLK218 plasmid pWLK-KPC | 35262 | *0.303* | *0.310* | *0.311* | *0.303* | 0.889 | 0.855 | 0.855 | 0.843 |

**Supplementary Figure 16:** Hierarchical clustering of plasmid sequences found in (a) *E. coli* and (b) *K. pneumoniae*, for subject 1674-T across various timepoints (based on Mash distance). Sequences with >95% identity were grouped together and a representative plasmid is shown in the leaves. (c) Table showing the proportion of each representative plasmid that is covered by contigs in various samples. Values in italics and grey indicate proportions that fall below 0.8 and were used to determine the presence, absence pattern shown in **Figure 3h**.

| Carbapenemase gene | *E. coli* | *K. pneumoniae* |
|---|---|---|
| KPC | 17 | 18 |
| OXA-48 | 42 | 31 |
| IMP | 6 | 0 |
| NDM | 8 | 13 |
| IMI | 0 | 1 |

**Supplementary Table 1:** Types of carbapenemase genes found in CPE *E. coli* and *K. pneumoniae* isolates.
